# Dissecting the loci underlying maturation timing in Atlantic salmon using haplotype and multi-SNP based association methods

**DOI:** 10.1101/2021.05.28.446127

**Authors:** Marion Sinclair-Waters, Torfinn Nome, Jing Wang, Sigbjørn Lien, Matthew P. Kent, Harald Sægrov, Bjørn Florø-Larsen, Geir H. Bolstad, Craig R. Primmer, Nicola J. Barson

## Abstract

Resolving the genetic architecture of fitness-related traits is key to understanding the evolution and maintenance of fitness variation. However, well-characterized genetic architectures of such traits in wild populations remain uncommon. In this study, we used haplotype-based and multi-SNP Bayesian association methods with sequencing data for 313 individuals from wild populations to further characterize known candidate regions for sea age at maturation in Atlantic salmon (*Salmo salar*). We detected an association at five loci (on chromosomes *ssa06*, *ssa09*, *ssa21*, and *ssa25*) out of 116 candidates previously identified in an aquaculture strain with maturation timing in wild Atlantic salmon. We found that at each of these five loci, variation explained by the locus was predominantly driven by a single SNP suggesting the genetic architecture of Atlantic salmon maturation includes multiple loci with simple, non-clustered alleles. This highlights the diversity of genetic architectures that can exist for fitness-related traits. Furthermore, this study provides a useful multi-SNP framework for future work using sequencing data to characterize genetic variation underlying phenotypes in wild populations.

## INTRODUCTION

Understanding the genetic processes underlying fitness variation is a fundamental goal in evolutionary biology. Identifying genetic variants that underlie fitness-related traits is therefore crucial, yet remains challenging. Substantial effort has been made to characterize the genetic architecture of traits – i.e. Are there few or many loci involved? Are loci effects small or large? How are loci distributed across the genome? And what are the allele frequencies at these loci [1–5]? It is generally assumed that in most cases single genetic variants translate into only small changes in complex traits, and therefore follow a polygenic [6,7] or an omnigenic [3,8] model of inheritance.

Among genome-wide association studies published to date, many complex traits appear to be polygenic [9]. Although polygenicity is widespread, an increasing number of examples of major effect loci exist, whereby one locus explains a large proportion of the phenotypic variation [10,11]. In some cases, major effect loci can contain multiple tightly linked genes, coined “supergenes”, where localized reduction in recombination is often caused by larger chromosomal rearrangements. For example, this phenomenon is known to underlie phenotypic variation observed among ruff (*Philomachus pugnax*) mating morphs [12,13], Atlantic cod (*Gadus morhua*) [14,15] and rainbow trout (*Oncorhynchus mykiss*) migratory ecotypes [16], and *Heliconius* butterfly wing-pattern morphs [17]. More recent work has found that major effect loci can exist alongside a polygenic background where loci with a variety of effect sizes underlie trait variation [18,19]. Such mixed genetic architectures may be pervasive, but currently remain undetected due to the large sample sizes required for detecting loci with smaller effects [19] and it is possible that additional examples are to be found with future higher-powered studies. Although studies aimed at resolving genotype-phenotype links are mounting, well-characterized genetic architectures of fitness-related traits, particularly in natural populations, are still uncommon.

While some trait-associated loci have been identified, such findings lead to other crucial questions: How have trait-locus associations arisen? Has the locus arisen through a single or multiple new mutations? Or alternatively, did the locus emerge via recombination that gave rise to new combinations of existing variants? Numerous studies from the past decade have shown that major effect loci involve the cumulative effects of multiple mutations, rather than a single mutation, thus highlighting the relevance of considering the latter scenarios. For example, Bickle et al. [20] found that ~60% of variation in female abdominal pigmentation in *Drosophila melanogaster* can be explained by sequence variation at the *bab* locus, but a GWAS (genome-wide association study) analyzing the same trait did not identify a single SNP in *bab* that passed the genome-wide significance threshold. Alleles consisting of multiple SNPs were associated with high proportions of the variation, whereas, single SNPs had only small effects and were therefore missed in the single-SNP GWAS. Additionally, Linnen at al. [11] and Kerdaffrec et al. [21] also identify multiple mutations within a confined region that have cumulative effects on colour traits in deer mice and seed dormancy in *Arabidopsis thaliana*, respectively. In natural populations with gene flow such as in Linnen et al. [11] and Kerdaffrec et al. [21], this is perhaps not unexpected as theory predicts that clustered and major effect loci will evolve under such scenarios [22,23]. Given these findings, examining extended sequence haplotypes containing multiple SNPs, rather than each SNP independently, is important [24]. This can be achieved by using alternative strategies that look at combined effects of variants, rather than single-SNP methods typically used in GWAS.

Here we investigate the genetic basis of Atlantic salmon (*Salmo salar*) sea age at maturity – the number of years spent in the marine environment before reaching maturity and returning to the natal river (freshwater) to reproduce. Age at maturity is an important life history trait affecting fitness traits such as survival, size at maturity and reproductive success [25,26]. Substantial variation in Atlantic salmon sea age at maturity is maintained due to a trade-off between mating success at spawning grounds and survival, whereby individuals that mature later are larger and have higher reproductive success on the spawning grounds, but lower survival and thus lower chance of reaching reproductive age. In contrast individuals that mature early are smaller and have lower reproductive success, but higher survival and thus higher chance of reaching reproductive age [27,28].

Variation in maturation timing in Atlantic salmon is highly heritable [19,29,30] and consequently there is substantial interest in understanding the underlying genetic architecture. A large-effect locus on chromosome 25 explaining up to 39% of the variation in sea age at maturity was found in wild European populations [10] and domesticated salmon [31]. The primary candidate gene underlying the association of this locus is *vgll3* due to its close proximity to the associated SNP variation [10,31,32] and its known function in other species. The *vgll3* gene encodes a transcription cofactor that, amongst other things, regulates adipogenesis [33] and is associated with variation in puberty timing in humans [34,35]. In addition to *vgll3*, Sinclair-Waters et al. [19] identified 119 other candidate genes for male maturation in a GWAS including >11,000 males from the same Atlantic salmon aquaculture strain. Two particularly strong associations between maturation timing were found on chromosome 9 in close proximity to *six6* and chromosome 25, *vgll3*. The association of *six6* was also found by Barson et al. [10] in wild Atlantic salmon, however, the signal disappeared after correction for population structure. Interestingly, the *six6* gene is also associated with age at maturity in two Pacific salmon species [36], humans [35] and cattle [37]. However, Barson et al. [10] focused solely on single-SNP associations via GWAS without considering the possible influence of combined variant effects.

Studies using sequencing data to examine variation associated with important fitness-related traits in wild populations are limited. However due to developments in sequencing technologies and bioinformatics, studies using this approach are likely to rise in number. We therefore aim to provide a useful and timely framework for characterizing genetic variation underlying phenotypes in wild populations in the future. Here, we focus on further characterizing the association between the loci identified in Sinclair-Waters et al. [15] and sea age at maturity in wild Atlantic salmon. We integrate re-sequencing data and phenotype information for 313 individuals from 53 wild population of Atlantic salmon with alternative GWAS strategies that consider the combined effects of variants, rather than single-SNP effects. This approach can provide better resolution of the variants that are potentially involved in controlling fitness-related traits such as maturation timing in Atlantic salmon.

## METHODS

### Study material

Whole genome sequencing data was obtained for 313 wild individuals collected from 53 Norwegian and Finnish populations spanning the Norwegian coast and to the Barents sea in the north (59°N - 71°N) (Supplementary Table S1) previously reported in Bertolotti et al. [38]. The 313-individual dataset includes populations belonging to both the Atlantic and Barents/White sea phylogeographic groups. These regions were studied in Barson et al. [10] using SNP-array data and a single SNP approach, therefore missing variants and potentially combined variant effects. Individuals were categorized into three maturation categories based on the number of years spent at sea prior to their first return migration to rivers for spawning: 1 (one year spent at sea), 2 (two years spent at sea), or 3 (three or more years spent at sea). Only five individuals had spent four years and were therefore combined with three-year fish for all analyses.

### SNP calling & filtering

Variant calling and the first round of filtering was done in a larger set of individuals described in Bertolotti et al. [38]. Raw Illumina reads were mapped to the Atlantic salmon genome (ICSASG_v2) [39] using *bcbio-nextgen v.1.1* [40]with the *bwa-mem aligner v.0.7.17* [41]. Genomic variation was identified using the Genome Analysis Toolkit (*GATK*) *v4.0.3.0*., following *GATK*’s best practice recommendations. *Picard v2.18.7* [42] was used to mark duplicates and *GATK* was used for joint calling [43]. Variants were annotated using *SNPeff v. 4.3* [44]. Variant call were further filtered with GATK’s variant filtration according to the following --*filterExpression*: “MQRankSum < −12.5 ∥ ReadPosRankSum < −8.0 ∥ QD < 2.0 ∥ FS > 60.0 ∥ (QD < 10.0 && AD[0:1] / (AD[0:1] + AD[0:0]) < 0.25 && ReadPosRankSum < 0.0) ∥ MQ < 30.0". SNPs were then filtered using *SNPable* procedure [45], where 100 bp kmers are mapped to reference genome (ICSASG_v2) using Burrows-Wheeler Aligner (*bwa aln*) [46], and only SNPs within regions with reads that uniquely map are retained. We then removed additional SNPs with *vcftools* using the following criteria: *--min-alleles 2, --max-alleles 2, --maf 0.0000000001, --max-missing 0.7, --remove-indels, --minGQ 10,* and –*minDP 4*. A subset 313 individuals from wild populations was then extracted from this larger dataset using *vcftools* [47]. This reduced dataset was used for all subsequent analyses.

### Principal component analysis

We produced a reduced SNP dataset by pruning one SNP from each SNP pair with a correlation coefficient (*r*^*2*^) greater than 0.2 within a 50 kb block using the *--indep-pairwise 50 10 0.2* function implemented in *PLINK v1.9* [48]. This yielded 403,540 SNPs to examine population structure using a principal component analysis, *smartpca,* implemented in the EIGENSOFT *v5* software [49].

### Data preparation

In this study, we focus on genomic regions containing the 116 candidate loci for age at maturity identified in Sinclair-Waters et al. [19]. We extracted SNP genotype data from 500 kb regions surrounding the 116 trait-associated SNPs identified in Sinclair-Waters et al. [19] using *vcftools’* [47] position filtering functions --*from-bp* and *--to-bp,* as well as allele filtering function *--mac 1* to keep only polymorphic sites. SNPs that were within 250 kb of an adjacent SNP were analyzed together by examining a region that extends 250 kb upstream of the first SNP to 250 kb downstream of the last SNP.

The current Atlantic salmon genome (ICSASG_v2) contains a known assembly error within the 500 kb region surrounding the known candidate loci *vgll3* [31]. A misplaced and misoriented scaffold currently placed downstream of *vgll3* belongs within a gap in the assembly just upstream of *vgll3* on ssa25. For this reason, we constructed a revised assembly for this chromosome. SNP calling was performed as described above. We then retained SNPs that had met the filtering criteria. A total of 8 candidate SNPs are located within regions of the genome that were moved. To find the position of these SNPs in the revised chromosome 25 sequence, we extracted 200 bp surrounding each of these SNPs from the current genome assembly (ICSASG_v2) using the *getfasta* function in *BEDTools* [50]. The 200 bp sequence was then blasted to the fixed assembly to determine the new position of each SNP using Blast’s *blastn* function [51]. Using the new SNP positions, SNP genotypes within a 500 kb region surrounding the moved candidate SNPs were extracted from the fixed dataset using *vcftools*.

### Association testing at candidate regions

We applied three association mapping methods to describe the genetic architecture underlying sea age at maturity at each of the candidate regions identified in Sinclair-Waters et al. [19]. First, a multi-SNP approach examining associations between phenotype and haplotypes was conducted using Bayesian linear regression implemented in *hapQTLv1.00* [52]. In this approach, a hidden Markov model is used to characterize haplotype structure and ancestry [53]. Haplotype sharing at each marker is then used to quantify genetic similarity among individuals. Haplotype associations are identified by testing for an association between genetic similarity at each marker and the phenotype [52]. Each of the extracted *vcf* files was converted to *bimbam* format using *PLINK 1.9* [54]. The resulting *bimbam* files were used as input for *hapQTL.* Second, single SNP associations were also identified using a Bayesian linear regression method implemented in *hapQTL* [55]. For all *hapQTL* association tests, sex and the six most significant principal components (see above) were included as covariates in the models. Each *hapQTL* run consisted of 2 EM runs (-e 2) with 40 steps (-w 40), 2 upper clusters (-C 2), 10 lower clusters (-c 10). Three replicate *hapQTL* runs were performed for each of the 116 selected regions. Based on recommendations from Jeffreys [56], Bayes factors greater than three were considered evidence for an association of either SNPs or haplotype with sea age at maturity phenotype.

Third, a multi-SNP approach aimed to estimate the number and identity of SNPs underlying trait variation at each candidate region using Bayesian Variable Selection regression implemented in *PiMASS* [55]. Due to computational restrictions, the *PiMASS* analysis was performed for only candidate regions that had a SNP or haplotype association with Bayes factor greater than 3. Prior to the *PiMASS* analysis, all missing genotypes were imputed in BIMBAM [55] as mean genotypes (-wmg) using default settings. Additionally, our phenotype values for sea age at maturity were adjusted to correct for confounding effects of sex and population structure by regressing the phenotype on sex and the six most significant principal components (see above) using the *lm* function in *R*. *PiMASS* was run with the residual phenotype values. We placed priors on the proportion of variance explained by SNP(s) (hmin = 0.001 and hmax = 0.999) and the number of SNPs in the model (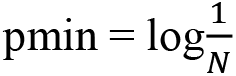 and 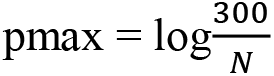, where N is the total number of SNPs). Each run consisted of a burn-in of 1000000 steps, followed by 2500000 steps where parameter values were recorded every 1000 steps. For each analysis, we examined the posterior inclusion probability for each SNP, the distribution of the number of included SNPs and the distribution of the proportions of variance explained per model. We also examined the path of estimated Bayes factors and parameter values (h, p, s) across all recorded iterations to check for convergence of runs.

To further assess whether more than one SNP in a candidate region was significantly associated with sea age at maturity, we regressed out the top-associated SNP from the residual phenotype values described above and reran *PiMASS* using the previously-used priors and settings. We then examined the posterior inclusion probability for each SNP, the distribution of the number of included SNPs, and the distribution of proportion of variance explained to determine whether there was evidence for multiple SNP associations within a given candidate region.

## RESULTS

### Principal component analysis

The first six principal components (PCs) calculated with the pruned SNP dataset explained 1.96%, 0.68%, 0.63%, 0.59%, 0.56% and 0.51% of the genetic variance, respectively (Supplementary Figure S1). These six PCs were included in subsequent association analyses to reflect population structure among samples.

### Associations identified with hapQTL

Single-SNP and haplotype association analyses with *hapQTL* revealed strong (Bayes factor > 3) association signals at 5 of the 116 candidate regions (Figure 1, Supplementary Figure S2). The strongest association observed within each region was with a single SNP, rather than an extended haplotype, suggesting a single mutation underlies the effect of each of these regions on maturation timing. However, exceptions occurred in the ssa09:24636574-25136574 and ssa25:28389273-28889273 regions, where second association signals were found upstream of the primary association signal and were most strongly linked to an extended haplotype. For instance, strong haplotype association scores (Bayes factor > 3) spanned a 26971 bp region (ssa09:24781742-24808713) containing an uncharacterized gene (LOC106610978) and *pcnx4.* In the ssa25:28389273-28889273 region, a strong haplotype signal was found within *edar* (Figure 1).

**Figure 1.**
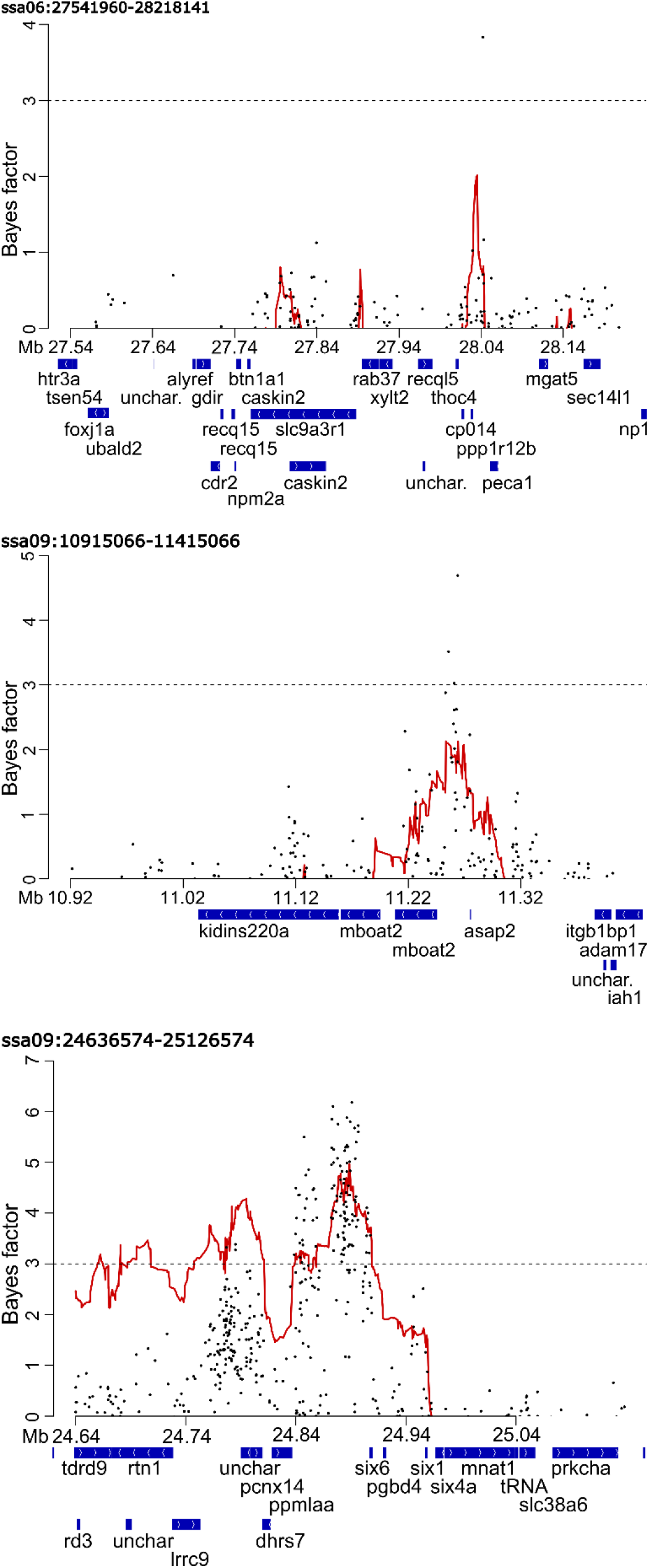
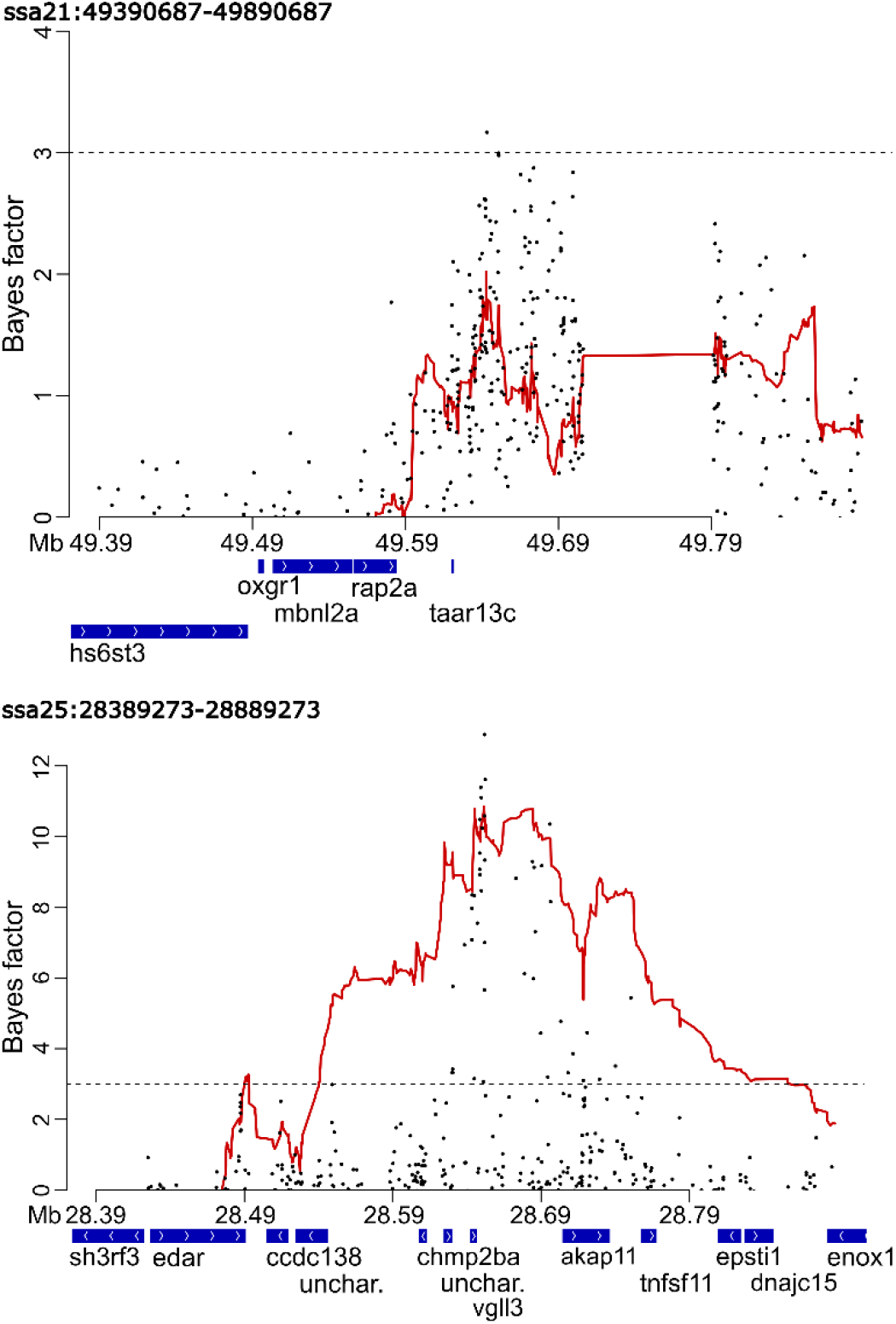
Plots displaying single SNP associations (black points) and haplotype associations (red line) scores from *hapQTL* for the five candidate regions with Bayes factors greater than 3. Y-axis shows the Bayes factor indicating the association strength. X-axis shows the position on the respective chromosomes.

We find differences in the location of the top-associated SNPs found here and those identified in Sinclair-Waters et al. [19]. For regions ssa06:27541960-28218141, ssa09:10915066-11415066 and ssa25:28389273-28889273, the top-associated SNP was located further upstream than in Sinclair-Waters et al. [19]. Contrastingly, the strongest associated SNPs within the regions ssa09:24636574-25136574 and ssa21:49390687-49890687 differed only slightly (<5000 bp) between studies (Table 1).

**Table 1.** Strongest association signals for each candidate region showing evidence of an association with sea age at maturity, the genes in closest proximity and association values from *hapQTL*. Top SNPs for each region from previous SNP-array study [19].

| Candidate region | Top signal | Closest gene | Bayes Factor | $-\log_{10}(P\text{-value})$ | Allele frequency | Top SNP(s)<sup>a</sup> | Candidate gene(s)<sup>a</sup> |
| --- | --- | --- | --- | --- | --- | --- | --- |
| ssa06:27541960-28218141 | 6:28045390 (SNP) | *pecam1*<br>(intron) | 3.835 | 5.107 | 0.320 | 6:27791960<br>6:27968141 | *slc9a3r1*<br>*recql5*<br>LOC106606978 |
| ssa09:10915066-11415066 | 9:11266848 (SNP) | *asap2a*<br>(upstream) | 4.696 | 5.434 | 0.074 | 9:11165066 | *mboat2* |
| ssa09:24636574-25136574 | 9:24888841 (SNP) | *six6*<br>(upstream) | 6.184 | 4.242 | 0.425 | 9:24886574 | *six6* |
| ssa21:49390687-49890687 | 21:49645222 (SNP) | *taar13c*<br>(upstream) | 3.172 | 4.649 | 0.464 | 21:49640687 | *taar13c* |
| ssa25:28389273-28889273 | 25: 28651640 (SNP)<br>[ICSASG_v2:<br>25:28669350] | *vgll3*<br>(downstream) | 12.893 | 6.406 | 0.358 | 25:28910202 | *vgll3* |

<sup>a</sup>From Sinclair-Waters et al. [19].

### Multi-SNP associations identified using PiMASS

Multi-SNP association analysis with *PiMASS* showed that at four of five candidate regions, a single-SNP model was most commonly used to explain variation in sea age at maturity. At one candidate region, ssa09:24636574-25136574, a multi-SNP model including two SNPs was most commonly used to explain variation in sea age at maturity. Median proportion of variance explained by each candidate region ranged between 4% and 19% (Figure 2, Table 2). However, when the top-associated SNP was regressed out from the phenotype values, no SNPs were selected to explain sea age at maturity for all five candidate regions. Additionally, post-regression median proportion of variance was substantially lower – ranging between 0% and 1% (Supplementary Figure S3, Table 2). This would suggest that sea age variation explained by each of these regions is largely driven by a single mutation. We observe no obvious trends in parameter values or Bayes factors, suggesting models converged and burn-in period was adequate (Supplementary Figure S4 &S5).

**Table 2.** *PiMASS* results prior to and after regression of top-associated SNP identified in the initial *PiMASS* analysis. These include the mode of the distribution of the number of SNPs and the median of the distribution of proportion of variance explained (PVE) for a model explaining sea age at maturity.

| Candidate region | Mode # of SNPs | Median PVE | Mode # of SNPs<br>(post-regression) | Median PVE<br>(post-regression) |
| --- | --- | --- | --- | --- |
| ssa06:27541960-28218141 | 1 | 0.05 | 0 | 0 |
| ssa09:10915066-11415066 | 1 | 0.07 | 0 | 0.01 |
| ssa09:24636574-25136574 | 2 | 0.09 | 0 | 0.01 |
| ssa21:49390687-49890687 | 1 | 0.04 | 0 | 0 |
| ssa25:28389273-28889273 | 1 | 0.19 | 0 | 0.01 |

**Figure 2.**
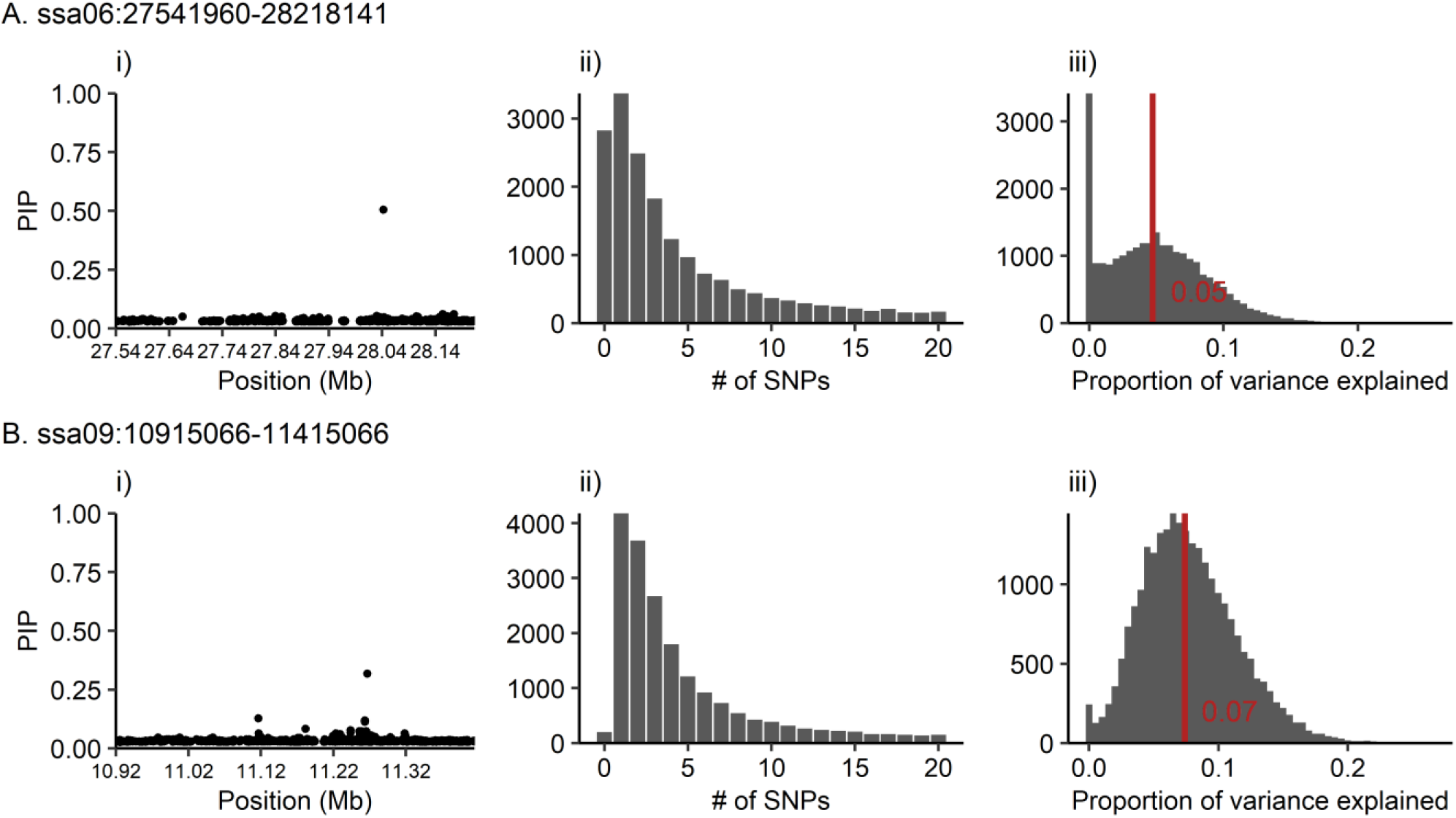
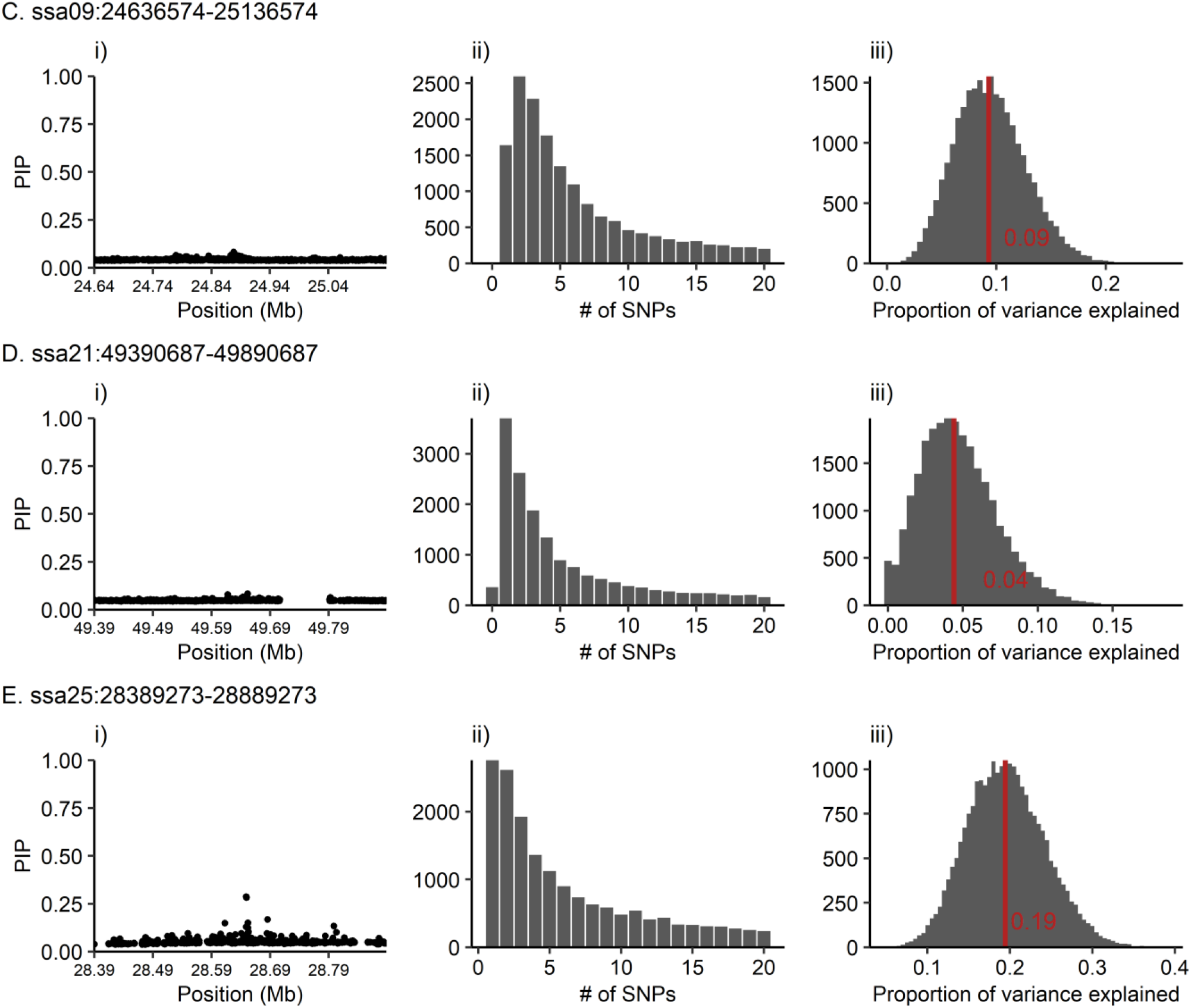
*PiMASS* results for each of the tested candidate regions: A. ssa06:27541960-28218141, B. ssa09:10915066-11415066 C. ssa09:24636574-25136574, D. ssa21:49390687-49890687, and E. ssa25:28389273-28889273. Plots display the following results for each candidate region: i) posterior inclusion probability (PIP) indicating the probability of a SNP being included in a model explaining sea age at maturity variation, ii) truncated distribution of the number of SNPs included in a model explaining sea age at maturity variation, and iii) distribution of proportion of variance explained per recorded iteration (2500). Red line indicates the median proportion of variance explained.

## DISCUSSION

Despite that combined effects of multiple variants at trait-associated loci are playing an important role in controlling fitness traits across a variety of species [11,20,21], our results indicate that sea age at maturation in Atlantic salmon is predominantly associated with single SNP variation at candidate regions. Using resequencing data to analyse 116 candidate loci and an analytical framework aimed at detecting multi-SNP associations, we find that single SNPs explain the variation in sea age at maturity in almost all cases. This work targeting candidate genes identified in aquaculture salmon strains suggests a mixed genetic architecture where a combination large-effect loci and smaller-effect loci also underlies age at maturity in wild Atlantic salmon populations. Two core loci, *vgll3* and *six6*, likely play a key role in determining age at maturity and additional smaller effect loci may be important for fine-tuning the trait across heterogeneous environments.

Theoretical modelling predicts that clustering of tightly linked adaptive mutations will occur under gene flow and selection in populations inhabiting spatially and/or temporally heterogeneous environments [22,23]. Although this seems to be a plausible scenario under which the genetic architecture of age at maturity has evolved in Atlantic salmon, our work suggests that the association in each of the candidate regions is driven by a single mutation. We cannot rule out, however, the possibility that the examined regions have pleiotropic effects and contain SNPs controlling other adaptive traits that have weak or no correlation with maturation timing. It is also possible that we did not have sufficient power to detect additional SNPs in these regions with small effects or with rare alleles. However, previous empirical studies have found few, but complex, loci with clusters of adaptive mutations [11,20,21], thus motivating our investigation of multi-SNP and haplotypic effects. Remington [24] also highlights the importance of distinguishing between allelic effects and single mutational effects when examining the genetic architecture of adaptive variation and its evolution. Our findings, however, suggest that alternative genetic architectures are feasible. One possible explanation could relate to the multiple whole genome duplication events that have occurred in Atlantic salmon and other salmonids [57]. The presence of multiple gene copies may impact the evolution of genetic architecture for traits such as age at maturity in Atlantic salmon. It is also possible that gene flow among Atlantic salmon populations is too restricted to neighbouring populations and/or strength of selection is insufficient for the establishment of linked mutations, as there is a rather specific balance of gene flow and selection required for clustered loci to arise [58]. Both an extension of models predicting genetic architecture and additional empirical studies – on a wider variety organisms and traits – are needed to evaluate the generality of particular architectures and to further understand the conditions under which they evolve.

We find additional evidence that a large-effect locus on ssa25, *vgll3,* largely underlies age at maturity in Atlantic salmon corroborating findings from a number of association studies on Atlantic salmon maturation [10,19,31,32,59]. The second strongest associated locus in this study is located in close proximity to *six6* on ssa09. This locus was previously found to be associated with early maturation in male farmed Atlantic salmon [19], with sea age at maturity in wild Atlantic salmon prior to population structure correction [10] and two species of Pacific salmon (Sockeye salmon and Steelhead trout) [36]. Additionally, we found another three loci associated with sea age at maturity: *pecam1, asap2aa* and *taar13c*. The handful of loci found here suggests that wild Atlantic salmon have a mixed genetic architecture where multiple loci, with a variety of effect sizes, control maturation timing – similar to what has been found in male farmed Atlantic salmon [19]. Knowledge of this mixed genetic architecture is highly relevant for how we predict the evolution of maturation timing in wild Atlantic salmon populations. A large body of work has shown the relevance of genetic architecture in determining evolutionary responses [60–68]. Recent works highlight the relevance of the genetic architecture underlying fitness traits when predicting a population’s response to environmental changes [69] and selective pressures such a fishing [70]. Future work elucidating how such mixed genetic architectures affect predicted evolution of traits, compared to that of omnigenic or polygenic architectures, will be valuable.

We find differences in locations of top-associated SNPs identified here and in Sinclair-Waters et al. [19]. This is not surprising given that we are examining sequence data that captures more SNP variation compared to SNP-array data used in Sinclair-Waters et al. [19]. Furthermore, we failed to find associations between sea age at maturity and many of the candidate regions identified in Sinclair-Waters et al. [19]. For example, several candidate regions on ssa03 and ssa04 displayed particularly strong association signals in aquaculture salmon, however, no signals at these regions were found here. Additionally, only one association peak at ssa06:27541960-28218141 was found here, whereas two independent associations within this region were found in aquaculture salmon [19]. Such differences may reflect changes in the genetic architecture of the trait evolving since the domestication of Atlantic salmon. Although, we would not expect large changes to occur given the domestication is relatively recent, just 10 to 15 generations ago [71]. Furthermore, this study is likely under-powered to detect all previously identified loci, particularly those with smaller effect sizes or rare alleles, due to smaller sample size. Additionally, there could be differences in genetic architecture among environments [72] and/or genotype by environment interactions giving rise to distinct genetic architectures in wild populations versus aquaculture strains.

We do not find strong evidence of multi-SNP associations at candidate loci examined in this study, however, we cannot yet disregard the utility of multi-SNP association methods for further resolving the genetic architecture of Atlantic salmon maturation. First, we do not examine the entire genome due to computational restrictions, rather, we focussed on 116 previously identified candidate regions. Second, the Atlantic salmon genome is highly complex [39] and therefore errors in the assembly that may be disruptive for haplotype-based analysis could exist. As new and improved versions of the Atlantic salmon genome are published, our ability to test for haplotypic associations will improve. Furthermore, in a few cases (ssa09:10915066-11415066, ssa09:24636574-25136574, ssa25:28389273-28889273) the *PiMASS* analyses post-regression of the top SNP selected no SNPs for a model explaining sea age at maturity variation, however, the median proportion of variance explained across all iterations was greater than zero. This may suggest that a weak signal was present, but was being missed due to insufficient power. Although this is largely speculative, it suggests that ruling out the possibility of multi-SNP associations at these particular candidate regions may be premature. Higher-powered studies (i.e. more individuals per population) may help to resolve this in the future.

In conclusion, our analytical framework, combining both single and multi-SNP association methods, reveals that single SNP variation is sufficient for explaining the association of previously identified candidate loci for Atlantic salmon maturation timing. Previous empirical and theoretical work have described trait-associated loci that have complex alleles with multiple variants, our findings therefore demonstrate the diversity of genetic architectures for fitness-related traits. Additional data, and a greater diversity of species and traits, will serve to better understand why this diversity of genetic architectures exists and how these particular genetic architectures evolve. The analytical framework used here will be a valuable resource for accomplishing this as individual-level resequencing data for wild species with phenotyped individuals becomes increasingly available.

## Acknowledgements

Funding was provided by Academy of Finland (grant numbers 307593, 302873 and 327255), the Research Council of Norway (NFR-275310 and NFR-275862) and a Natural Sciences and Engineering Research Council of Canada postgraduate scholarship. Wild Atlantic salmon genome sequencing was funded by the Research Council of Norway (The Aqua Genome project; ref: 221734). We would like to acknowledge Terese Andersstuen, Dr Mariann Árnyasi and Hanna Hellerud Hansen from CIGENE for their work in organising the sequencing of samples. We thank Gunnel Østborg (NINA), Kurt Urdal (Rådgivende Biologer) and Natural Resources Institute Finland (LUKE) for their work collecting phenotype data. We also acknowledge the Aqua Genome project for providing access to data prior to public release. The Orion Computing Cluster at CIGENE-NMBU and CSC – IT Center for Science, Finland are acknowledged for computational resources. Storage resources were provided by the Norwegian National Infrastructure for Research Data (NIRD, project NS9055K). Phenotype data was provided by the Norwegian Institute for Nature Research (NINA).

## Data availability

Genome re-sequencing data for individuals used in this study are available in the European Nucleotide Archive (ENA) or NCBI with the project accession code PRJEB38061 [38].

## Contributions

CRP, NJB, MSW conceived the study. TN developed the variant calling workflow and constructed the fixed assembly of *ssa25*. JW developed the variant filtering criteria. MSW performed all downstream analyses with input from NJB. MPK played key role in generating whole genome sequencing data. SL led the whole genome sequencing work as part of the AquaGenome project. HS, GHB, BFL, CRP coordinated Atlantic salmon sampling and provided phenotypic information. MSW, CRP, NJB drafted the manuscript. All authors commented on and approved the final manuscript.

