## Supplementary Table for "Dissecting the loci underlying maturation timing in Atlantic salmon using haplotype and multi-SNP based association methods"

Supplementary Table S1. List of river names, river ID, latitude and longitude coordinates, and number of individuals (N) collected at each location.

| River | River ID | Latitude | Longitude | N |
|---|---|---|---|---|
| Alta | 212.Z | 69.968 | 23.375 | 10 |
| Årgårdsvassdraget | 138.Z | 64.312 | 11.223 | 10 |
| Årøy | 077.Z | 61.268 | 7.1671 | 2 |
| Beiarvassdraget | 161.Z | 67.028 | 14.579 | 2 |
| Børselva in Porsanger | 225.Z | 70.312 | 25.539 | 2 |
| Daleelva (Høyangervassdraget) | 079.Z | 61.219 | 6.0748 | 2 |
| Driva | 109.Z | 62.67606 | 8.550612 | 8 |
| Eidfjordvassdraget | 050.Z | 60.466 | 7.0718 | 2 |
| Eira | 104.Z | 62.678 | 8.1196 | 2 |
| Elvegårdselva (Bjerkvik) | 175.Z | 68.546 | 17.562 | 1 |
| Enningdalselva | 001.1Z | 58.981 | 11.474 | 10 |
| Ervikelva | 091.3Z | 68.8218 | 16.48854 | 10 |
| Etneelva | 041.Z | 59.67 | 5.9416 | 2 |
| Flåmselva | 072.2Z | 60.8651 | 7.1186 | 10 |
| Flekkeelva | 082.Z | 61.31 | 5.3445 | 2 |
| Forsåvassdraget | 172.Z | 68.151 | 16.116 | 2 |
| Gaula in Sør-Trøndelag | 122.Z | 63.341 | 10.236 | 2 |
| Gloppenelva | 087.Z | 61.768 | 6.2 | 2 |
| Homla | 123.4Z | 63.413 | 10.804 | 10 |
| Jølstra | 084.Z | 61.455 | 5.8434 | 3 |
| Komagelva | 239.Z | 70.242 | 30.522 | 10 |
| Lærdalselva | 073.Z | 61.102 | 7.4725 | 2 |
| Tana (Laksjohka) | 234.Z | 70.059 | 27.562 | 3 |
| Lakselva in Porsanger | 224.Z | 70.078 | 24.927 | 3 |
| Langfjordelva | 233.Z | 70.66914 | 27.83905 | 10 |
| Laukhellevassdraget (Lakselva from Trollbuvatnet) | 194.Z | 69.227 | 17.849 | 10 |
| Loneelva in Osterøy | 060.4Z | 60.52 | 5.5011 | 2 |
| Målselvvassdraget | 196.Z | 69.264 | 18.51 | 9 |
| Tana (Maskejohka) | 234.Z | 70.285 | 28.163 | 3 |
| Namsen (hele vassdraget) | 139.Z | 64.464 | 11.682 | 5 |
| Namsen (Fiskumfoss/Tørris) | 139Z | 64.60584 | 12.54821 | 5 |
| Nausta | 084.7Z | 61.506 | 5.7197 | 10 |
| Neiden (Näätämöjoki in Finnish) | 244.Z | 69.70143 | 29.52523 | 10 |
| Numedalslågen | 015.Z | 59.06 | 10.071 | 10 |
| Orkla | 121.Z | 63.30663 | 9.82683 | 10 |
| Oselva in Os | 085.Z | 60.186 | 5.4723 | 2 |
| Oselva in Osen | 055.7Z | 61.55063 | 5.413458 | 10 |
| Reipåga | 160.43Z | 66.908 | 13.632 | 10 |
| Repparfjordelva | 213.Z | 70.445 | 24.328 | 10 |
| Risfjordvassdraget | 231.8Z | 70.978 | 28.171 | 3 |
| Roksdalsvassdraget | 186.2Z | 69.05 | 15.869 | 2 |
| Ryggelva | 087.1Z | 61.779 | 6.1249 | 10 |
| Saltdalsvassdraget | 163.Z | 67.098 | 15.419 | 10 |
| Sandfjordelva in Gamvik | 231.7Z | 71.049 | 28.057 | 3 |

| | | | | |
|---|---|---|---|---|
| Skienselva | 016.Z | 59.135 | 9.6301 | 2 |
| Skipsfjordvassdraget | 202.11Z | 70.158 | 19.797 | 2 |
| Suldalslågen | 036.Z | 59.48 | 6.2506 | 10 |
| Surna | 112.Z | 62.971 | 8.6624 | 2 |
| Sylteelva in Fræna | 107.3Z | 62.838 | 7.2096 | 2 |
| Teno (lower-Utsjoki) | 234.Z | 69.909 | 27.0285 | 14 |
| Vestre Jakobselv | 240.Z | 70.108 | 29.327 | 3 |
| Vigda | 122.2Z | 63.312 | 10.182 | 10 |
| Vikedalselva in Vindafjord | 038.Z | 59.496 | 5.8972 | 10 |
| Vorma | 035.3Z | 59.271 | 6.3322 | 2 |
