## Supplementary Figures for "Dissecting the loci underlying maturation timing in Atlantic salmon using haplotype and multi-SNP based association methods"

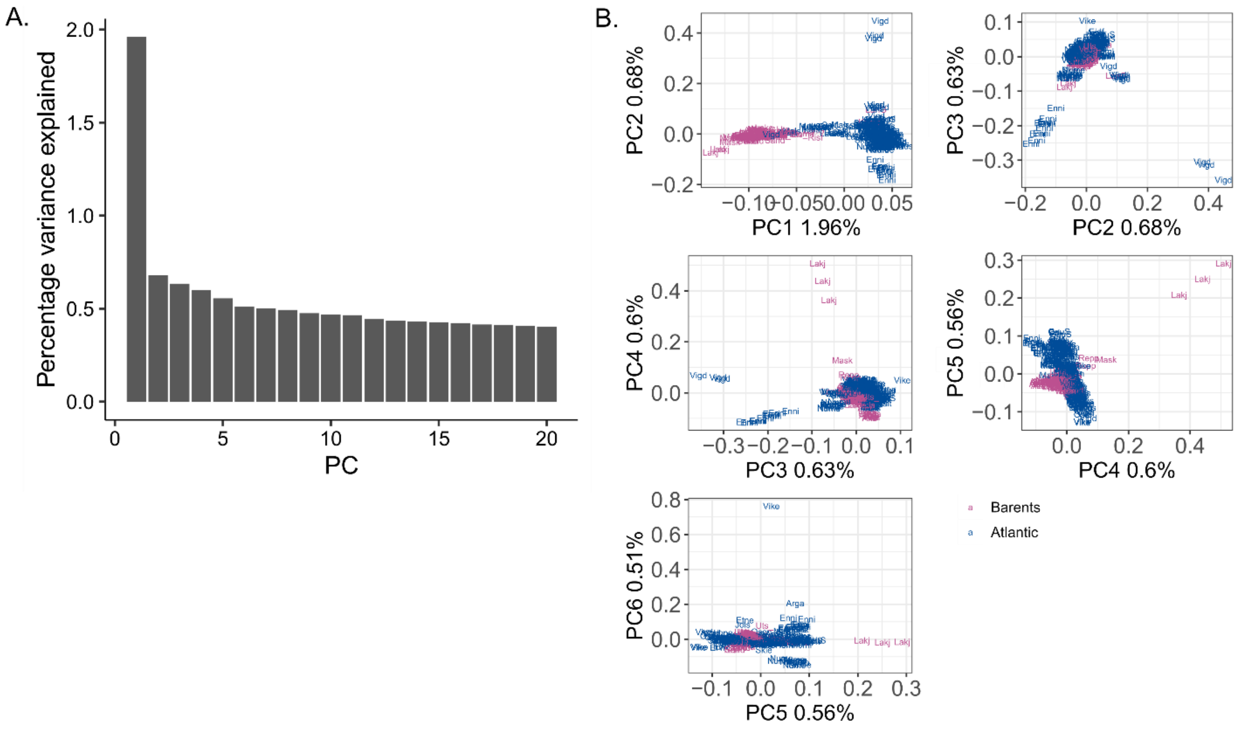


Supplementary Figure S1. A) Percentage of variance explained by the first 20 principal components. B) PCA plots displaying the first six principal components.

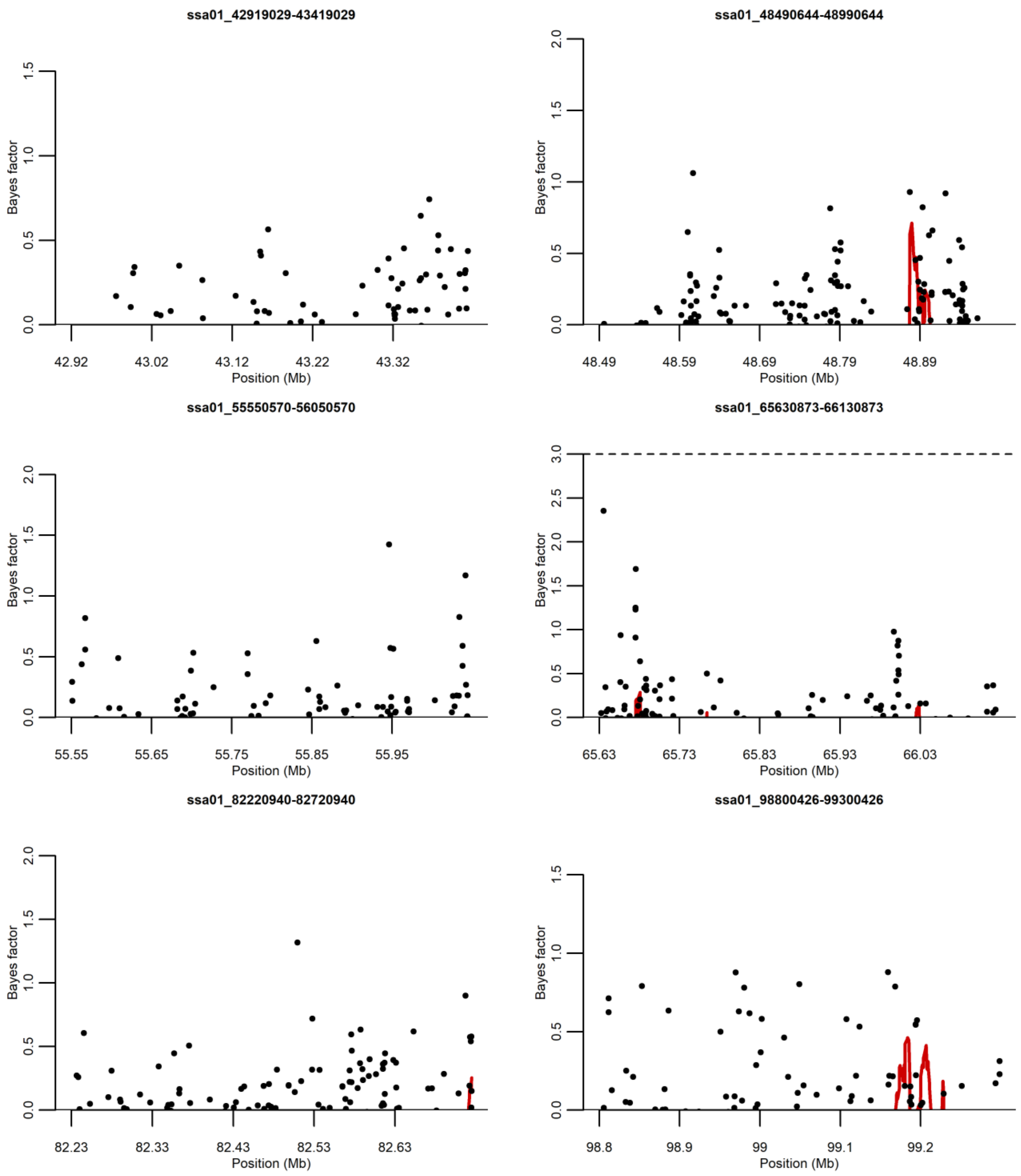
ssa01_42919029-43419029
Bayes factor
0.0
0.5
1.0
1.5
42.92
43.02
43.12
43.22
43.32
Position (Mb)
ssa01_48490644-48990644
Bayes factor
0.0
0.5
1.0
1.5
2.0
48.49
48.59
48.69
48.79
48.89
Position (Mb)
ssa01_55550570-56050570
Bayes factor
0.0
0.5
1.0
1.5
2.0
55.55
55.65
55.75
55.85
55.95
Position (Mb)
ssa01_65630873-66130873
Bayes factor
0.0
0.5
1.0
1.5
2.0
2.5
3.0
65.63
65.73
65.83
65.93
66.03
Position (Mb)
ssa01_82220940-82720940
Bayes factor
0.0
0.5
1.0
1.5
2.0
82.23
82.33
82.43
82.53
82.63
Position (Mb)
ssa01_98800426-99300426
Bayes factor
0.0
0.5
1.0
1.5
98.8
98.9
99
99.1
99.2
Position (Mb)

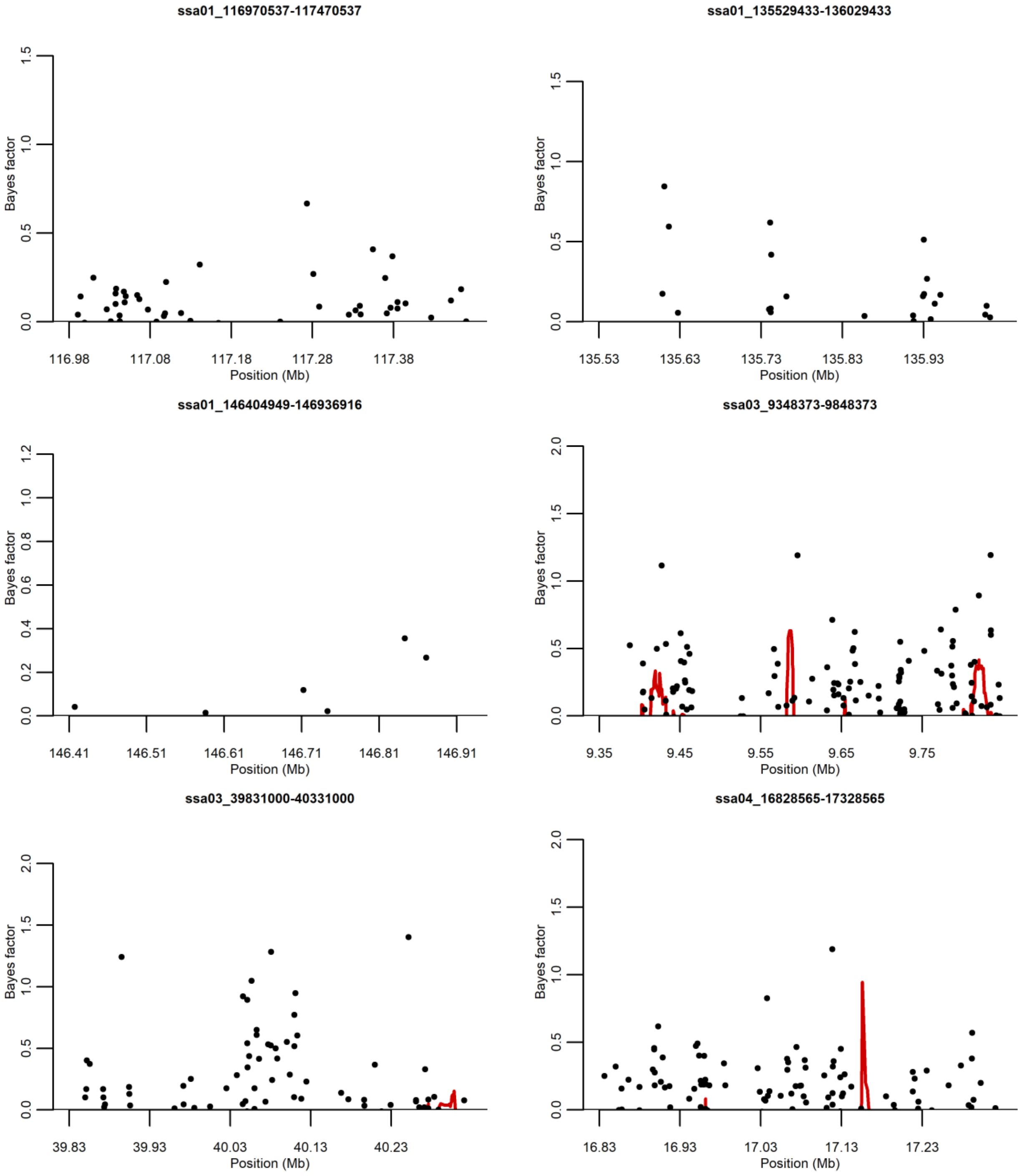
ssa01_116970537-117470537
Bayes factor
0.0
0.5
1.0
1.5
116.98
117.08
117.18
117.28
117.38
Position (Mb)
ssa01_135529433-136029433
Bayes factor
0.0
0.5
1.0
1.5
135.53
135.63
135.73
135.83
135.93
Position (Mb)
ssa01_146404949-146936916
Bayes factor
0.0
0.2
0.4
0.6
0.8
1.0
1.2
146.41
146.51
146.61
146.71
146.81
146.91
Position (Mb)
ssa03_9348373-9848373
Bayes factor
0.0
0.5
1.0
1.5
2.0
9.35
9.45
9.55
9.65
9.75
Position (Mb)
ssa03_39831000-40331000
Bayes factor
0.0
0.5
1.0
1.5
2.0
39.83
39.93
40.03
40.13
40.23
Position (Mb)
ssa04_16828565-17328565
Bayes factor
0.0
0.5
1.0
1.5
2.0
16.83
16.93
17.03
17.13
17.23
Position (Mb)

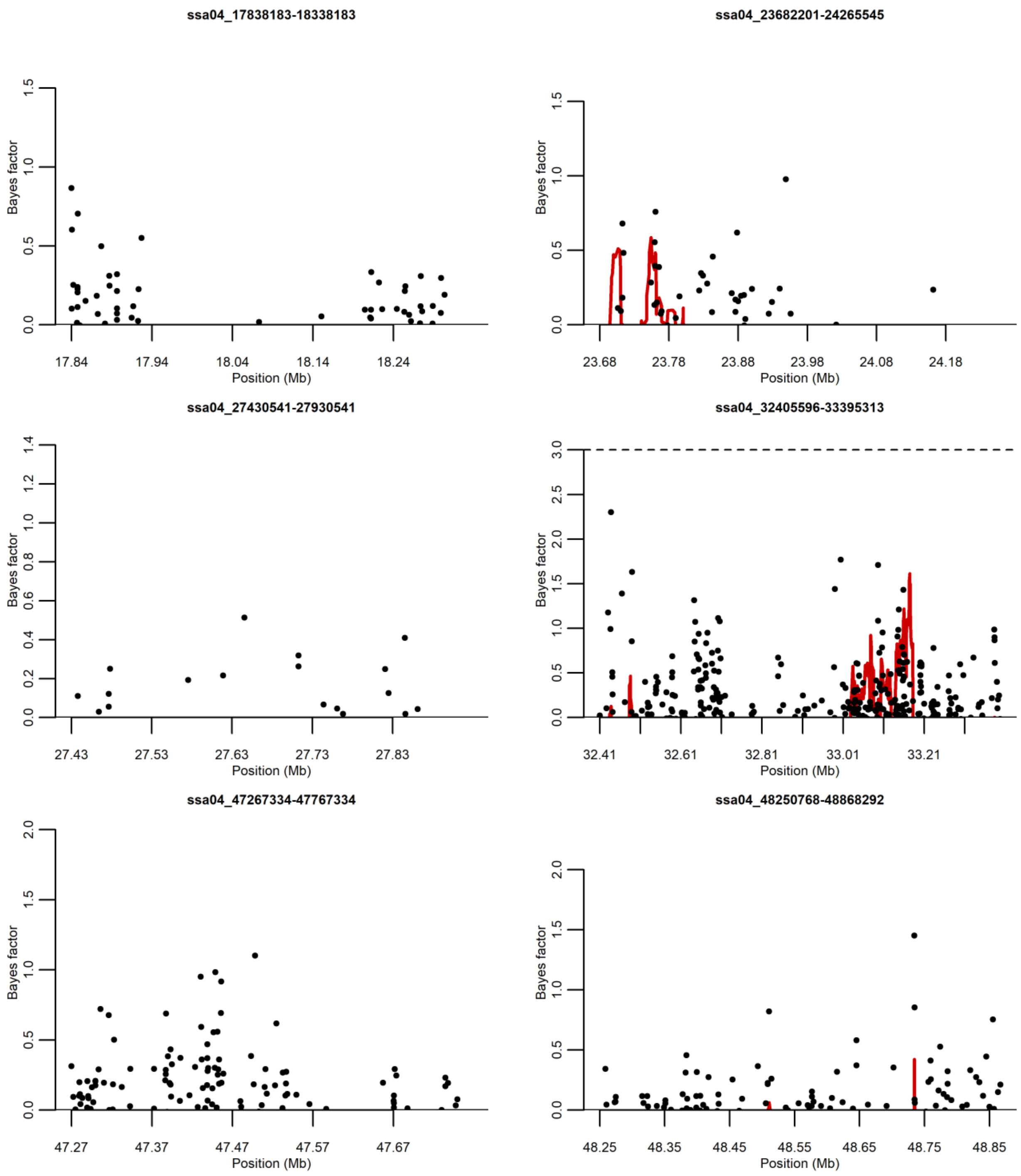

ssa04_17838183-18338183
ssa04_23682201-24265545
ssa04_27430541-27930541
ssa04_32405596-33395313
ssa04_47267334-47767334
ssa04_48250768-48868292
Bayes factor
Position (Mb)

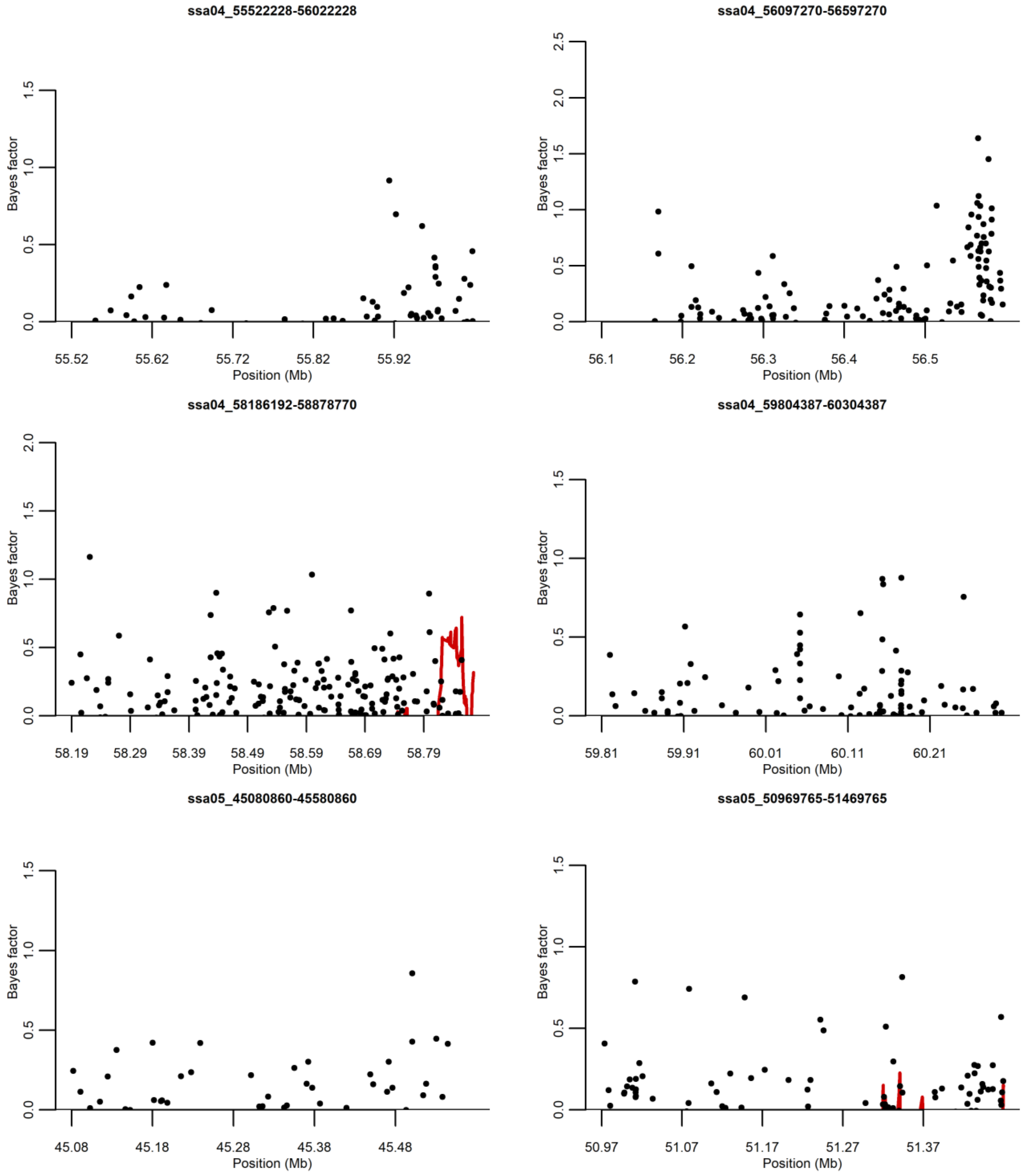
ssa04_55522228-56022228
Bayes factor
0.0
0.5
1.0
1.5
55.52
55.62
55.72
55.82
55.92
Position (Mb)
ssa04_56097270-56597270
Bayes factor
0.0
0.5
1.0
1.5
2.0
2.5
56.1
56.2
56.3
56.4
56.5
Position (Mb)
ssa04_58186192-58878770
Bayes factor
0.0
0.5
1.0
1.5
2.0
58.19
58.29
58.39
58.49
58.59
58.69
58.79
Position (Mb)
ssa04_59804387-60304387
Bayes factor
0.0
0.5
1.0
1.5
59.81
59.91
60.01
60.11
60.21
Position (Mb)
ssa05_45080860-45580860
Bayes factor
0.0
0.5
1.0
1.5
45.08
45.18
45.28
45.38
45.48
Position (Mb)
ssa05_50969765-51469765
Bayes factor
0.0
0.5
1.0
1.5
50.97
51.07
51.17
51.27
51.37
Position (Mb)

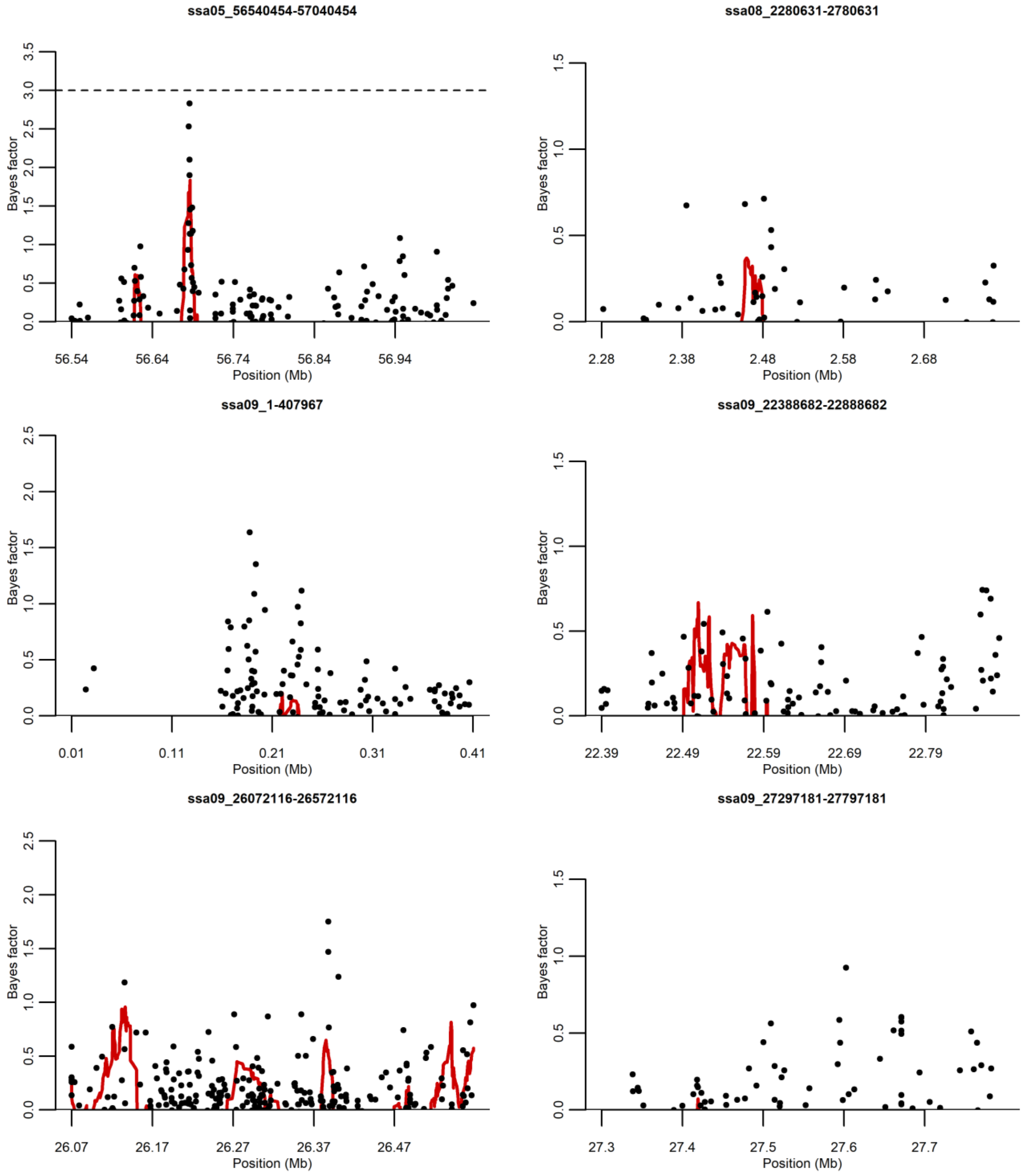
ssa05_56540454-57040454
Bayes factor
Position (Mb)
56.54
56.64
56.74
56.84
56.94
ssa08_2280631-2780631
Bayes factor
Position (Mb)
2.28
2.38
2.48
2.58
2.68
ssa09_1-407967
Bayes factor
Position (Mb)
0.01
0.11
0.21
0.31
0.41
ssa09_22388682-22888682
Bayes factor
Position (Mb)
22.39
22.49
22.59
22.69
22.79
ssa09_26072116-26572116
Bayes factor
Position (Mb)
26.07
26.17
26.27
26.37
26.47
ssa09_27297181-27797181
Bayes factor
Position (Mb)
27.3
27.4
27.5
27.6
27.7

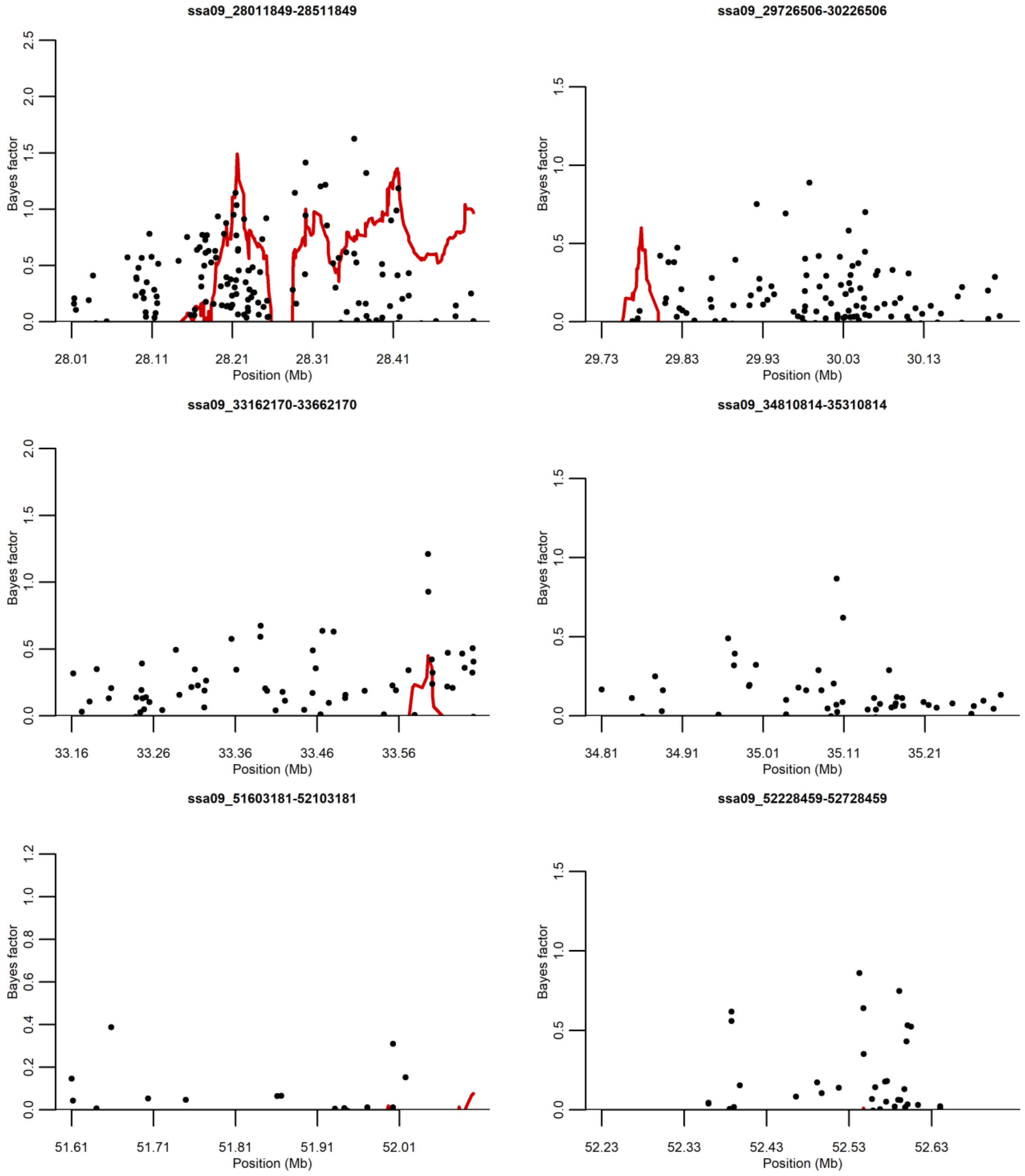
ssa09_28011849-28511849
ssa09_29726506-30226506
ssa09_33162170-33662170
ssa09_34810814-35310814
ssa09_51603181-52103181
ssa09_52228459-52728459
Bayes factor
Position (Mb)

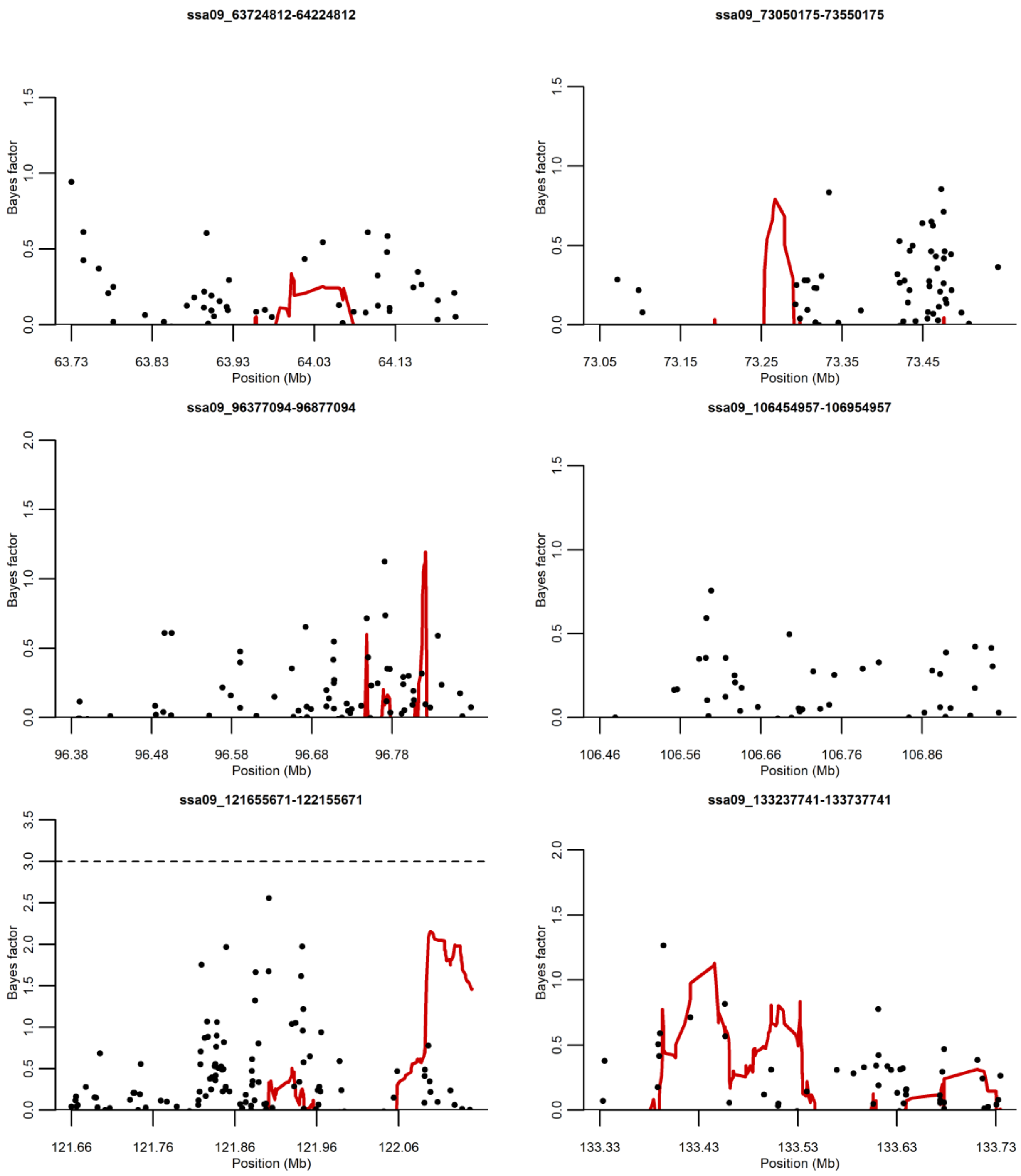

ssa09_63724812-64224812
Bayes factor
Position (Mb)
63.73
63.83
63.93
64.03
64.13
ssa09_73050175-73550175
Bayes factor
Position (Mb)
73.05
73.15
73.25
73.35
73.45
ssa09_96377094-96877094
Bayes factor
Position (Mb)
96.38
96.48
96.58
96.68
96.78
ssa09_106454957-106954957
Bayes factor
Position (Mb)
106.46
106.56
106.66
106.76
106.86
ssa09_121655671-122155671
Bayes factor
Position (Mb)
121.66
121.76
121.86
121.96
122.06
ssa09_133237741-133737741
Bayes factor
Position (Mb)
133.33
133.43
133.53
133.63
133.73

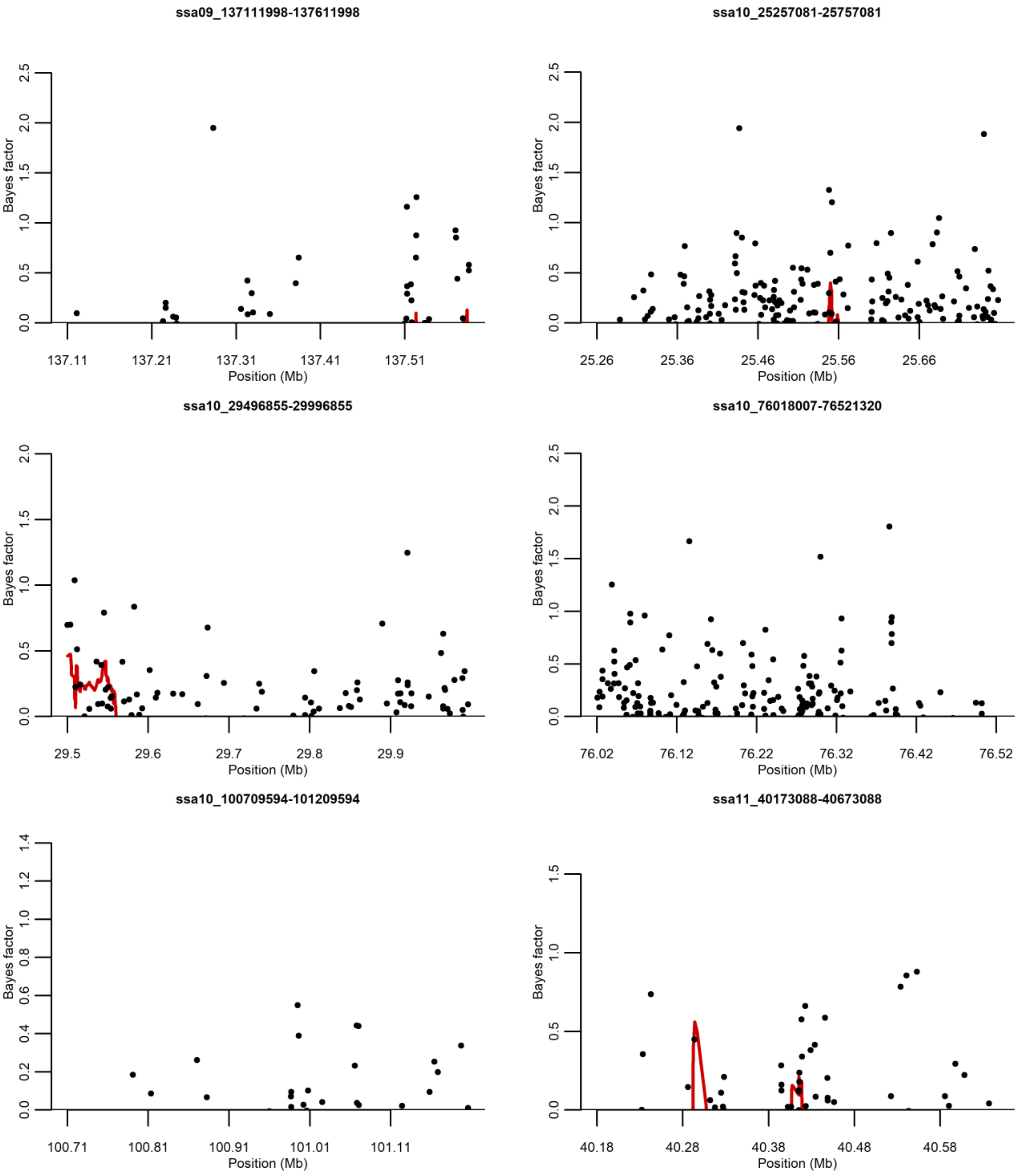
ssa09_137111998-137611998
Bayes factor
Position (Mb)
137.11
137.21
137.31
137.41
137.51
ssa10_25257081-25757081
Bayes factor
Position (Mb)
25.26
25.36
25.46
25.56
25.66
ssa10_29496855-29996855
Bayes factor
Position (Mb)
29.5
29.6
29.7
29.8
29.9
ssa10_76018007-76521320
Bayes factor
Position (Mb)
76.02
76.12
76.22
76.32
76.42
76.52
ssa10_100709594-101209594
Bayes factor
Position (Mb)
100.71
100.81
100.91
101.01
101.11
ssa11_40173088-40673088
Bayes factor
Position (Mb)
40.18
40.28
40.38
40.48
40.58

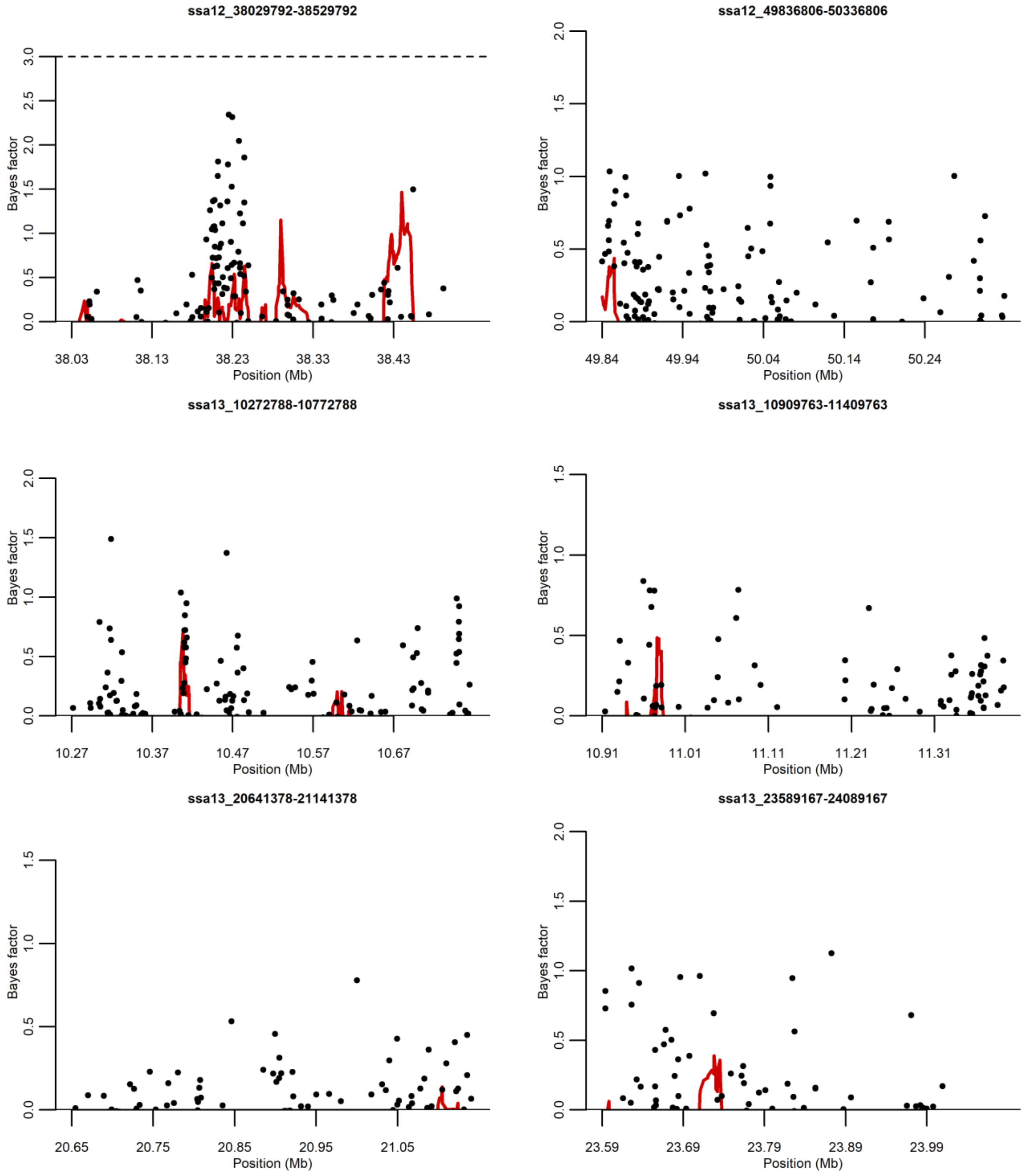
ssa12_38029792-38529792
Bayes factor
Position (Mb)
38.03
38.13
38.23
38.33
38.43
ssa12_49836806-50336806
49.84
49.94
50.04
50.14
50.24
ssa13_10272788-10772788
10.27
10.37
10.47
10.57
10.67
ssa13_10909763-11409763
10.91
11.01
11.11
11.21
11.31
ssa13_20641378-21141378
20.65
20.75
20.85
20.95
21.05
ssa13_23589167-24089167
23.59
23.69
23.79
23.89
23.99

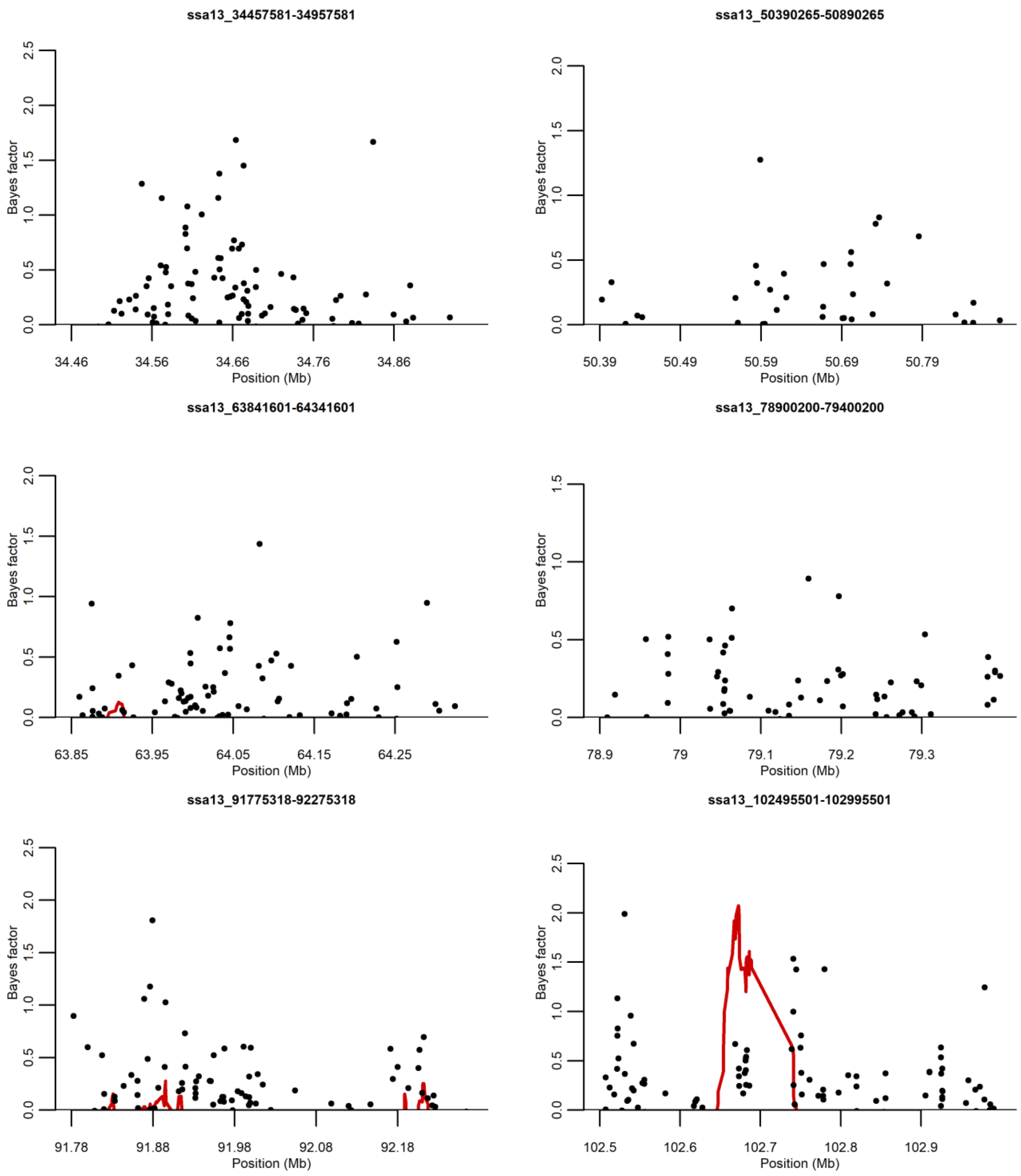
ssa13_34457581-34957581
ssa13_50390265-50890265
ssa13_63841601-64341601
ssa13_78900200-79400200
ssa13_91775318-92275318
ssa13_102495501-102995501
Bayes factor
Position (Mb)
34.46
34.56
34.66
34.76
34.86
50.39
50.49
50.59
50.69
50.79
63.85
63.95
64.05
64.15
64.25
78.9
79
79.1
79.2
79.3
91.78
91.88
91.98
92.08
92.18
102.5
102.6
102.7
102.8
102.9

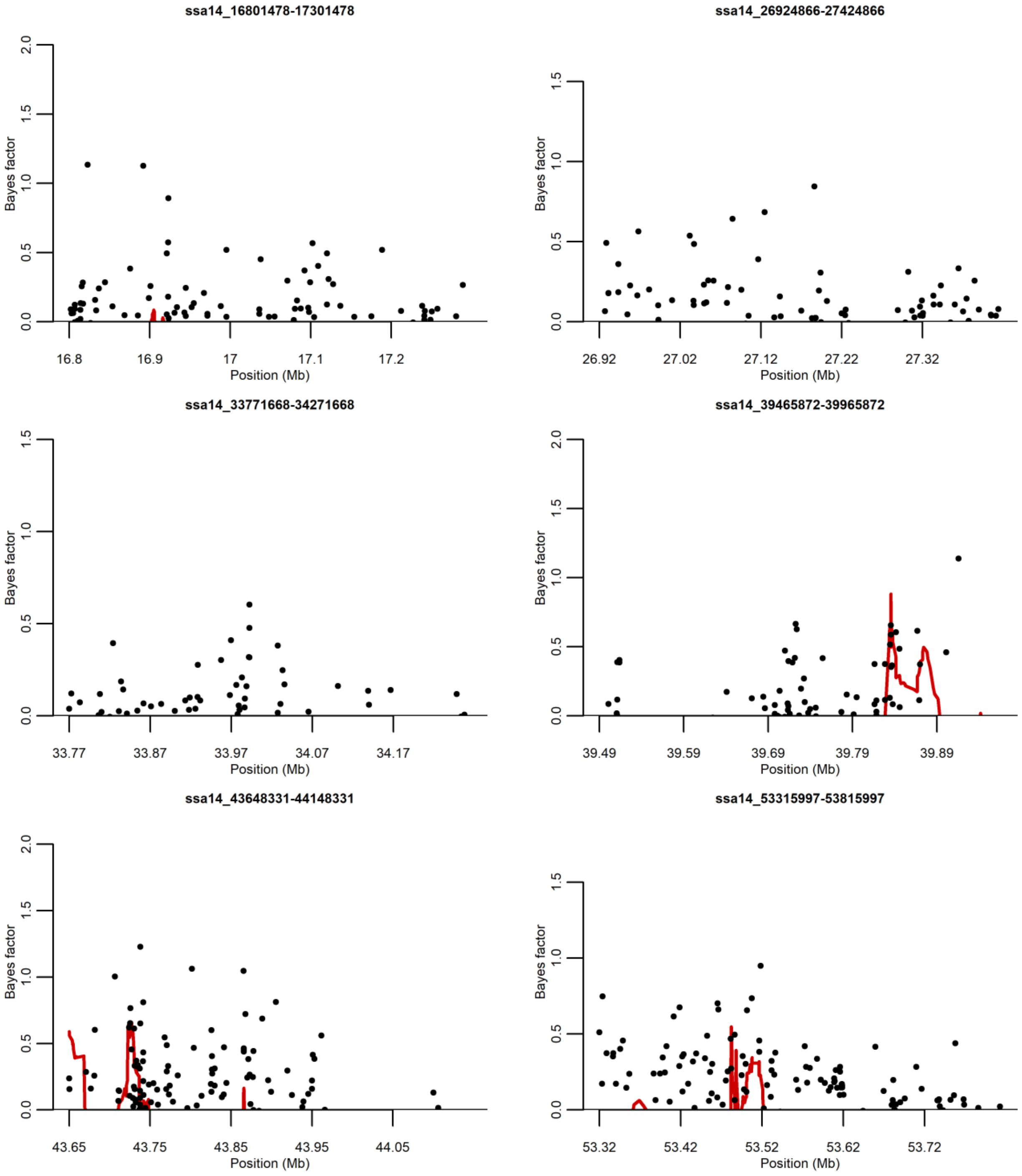

ssa14_16801478-17301478
Bayes factor
0.0
0.5
1.0
1.5
2.0
16.8
16.9
17
17.1
17.2
Position (Mb)
ssa14_26924866-27424866
Bayes factor
0.0
0.5
1.0
1.5
26.92
27.02
27.12
27.22
27.32
Position (Mb)
ssa14_33771668-34271668
Bayes factor
0.0
0.5
1.0
1.5
33.77
33.87
33.97
34.07
34.17
Position (Mb)
ssa14_39465872-39965872
Bayes factor
0.0
0.5
1.0
1.5
2.0
39.49
39.59
39.69
39.79
39.89
Position (Mb)
ssa14_43648331-44148331
Bayes factor
0.0
0.5
1.0
1.5
2.0
43.65
43.75
43.85
43.95
44.05
Position (Mb)
ssa14_53315997-53815997
Bayes factor
0.0
0.5
1.0
1.5
53.32
53.42
53.52
53.62
53.72
Position (Mb)

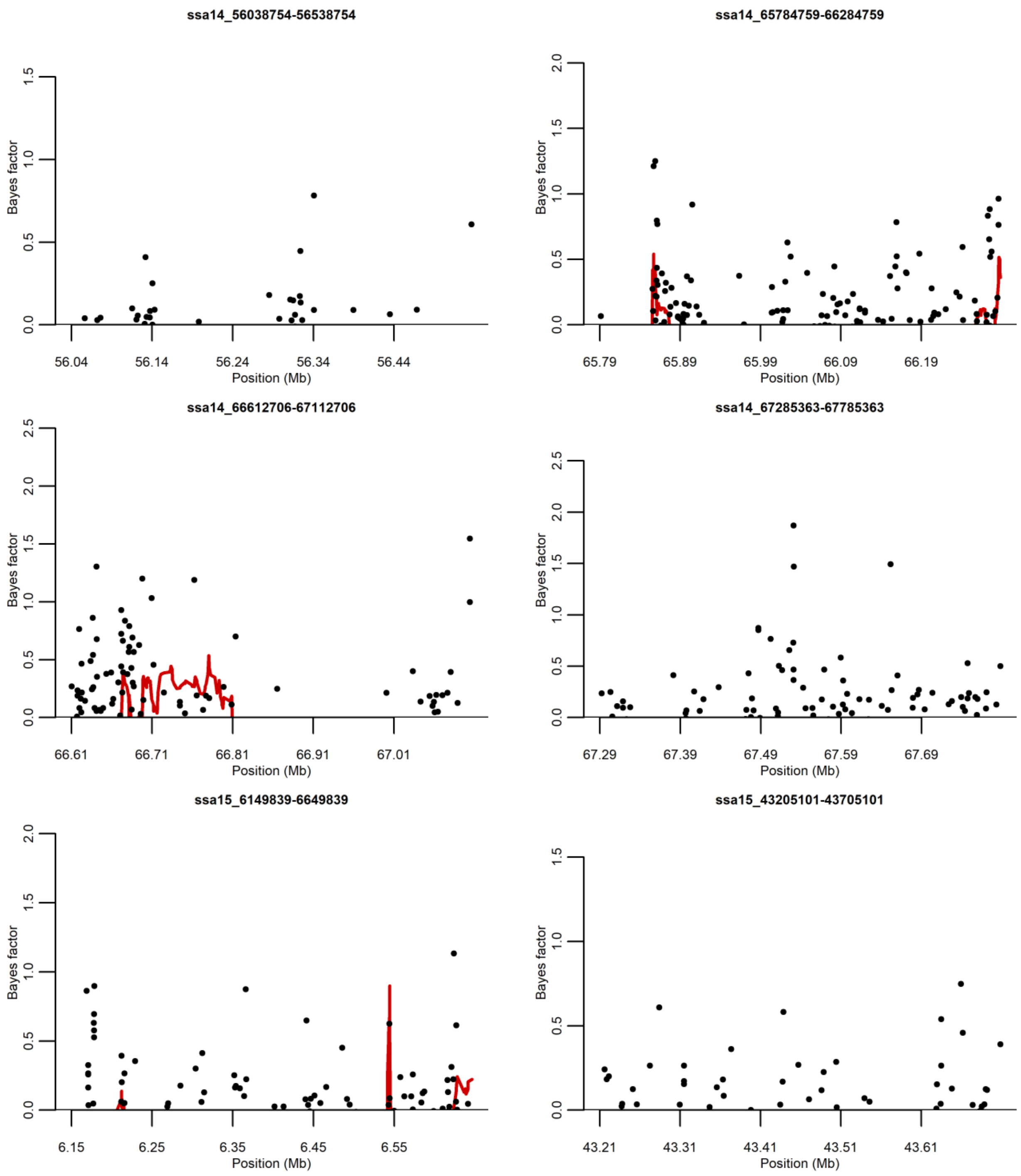
ssa14_56038754-56538754
Bayes factor
Position (Mb)
56.04
56.14
56.24
56.34
56.44
ssa14_65784759-66284759
Bayes factor
Position (Mb)
65.79
65.89
65.99
66.09
66.19
ssa14_66612706-67112706
Bayes factor
Position (Mb)
66.61
66.71
66.81
66.91
67.01
ssa14_67285363-67785363
Bayes factor
Position (Mb)
67.29
67.39
67.49
67.59
67.69
ssa15_6149839-6649839
Bayes factor
Position (Mb)
6.15
6.25
6.35
6.45
6.55
ssa15_43205101-43705101
Bayes factor
Position (Mb)
43.21
43.31
43.41
43.51
43.61

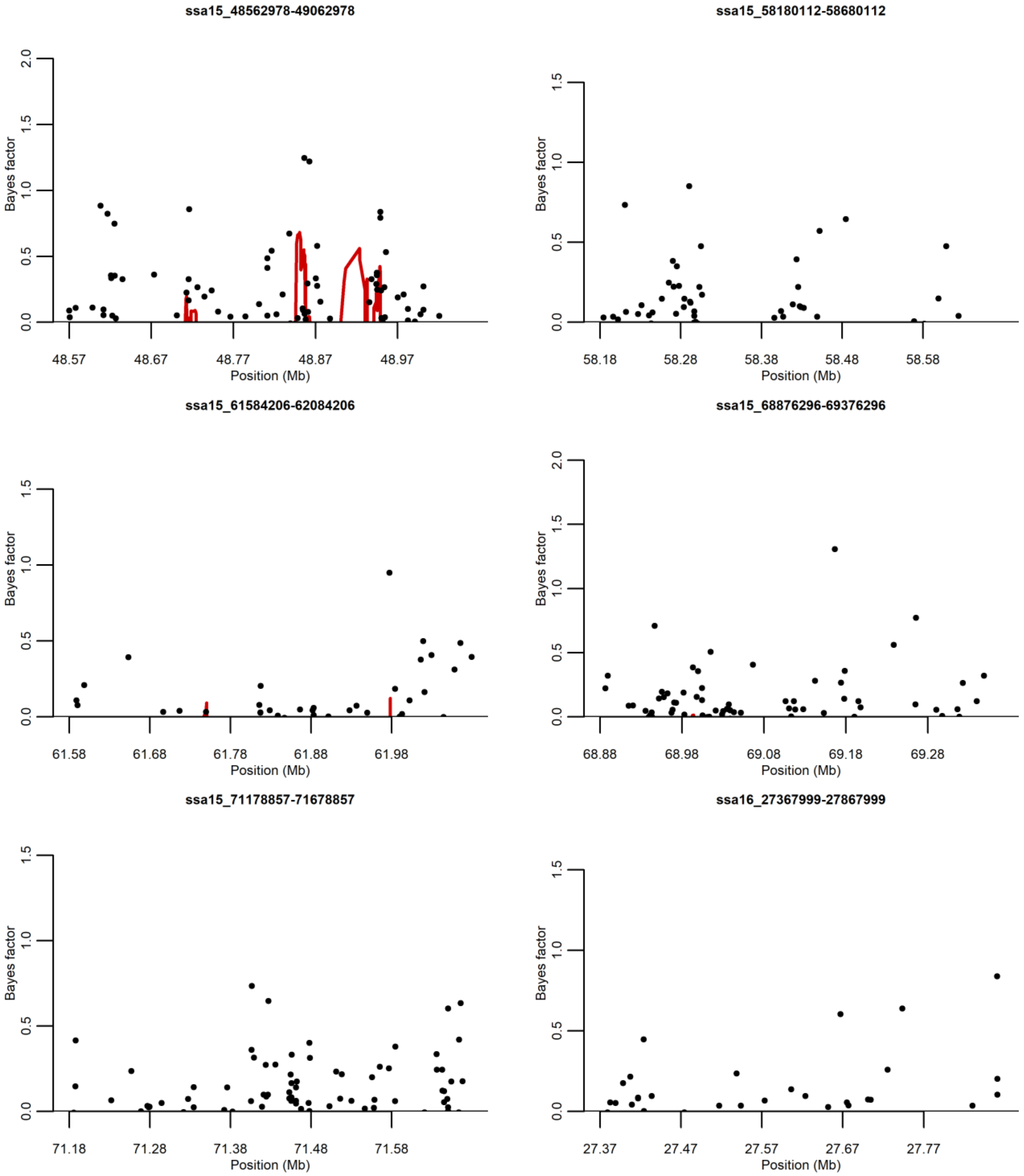
ssa15_48562978-49062978
ssa15_58180112-58680112
ssa15_61584206-62084206
ssa15_68876296-69376296
ssa15_71178857-71678857
ssa16_27367999-27867999
Bayes factor
Position (Mb)
0.0
0.5
1.0
1.5
2.0
48.57
48.67
48.77
48.87
48.97
58.18
58.28
58.38
58.48
58.58
61.58
61.68
61.78
61.88
61.98
68.88
68.98
69.08
69.18
69.28
71.18
71.28
71.38
71.48
71.58
27.37
27.47
27.57
27.67
27.77

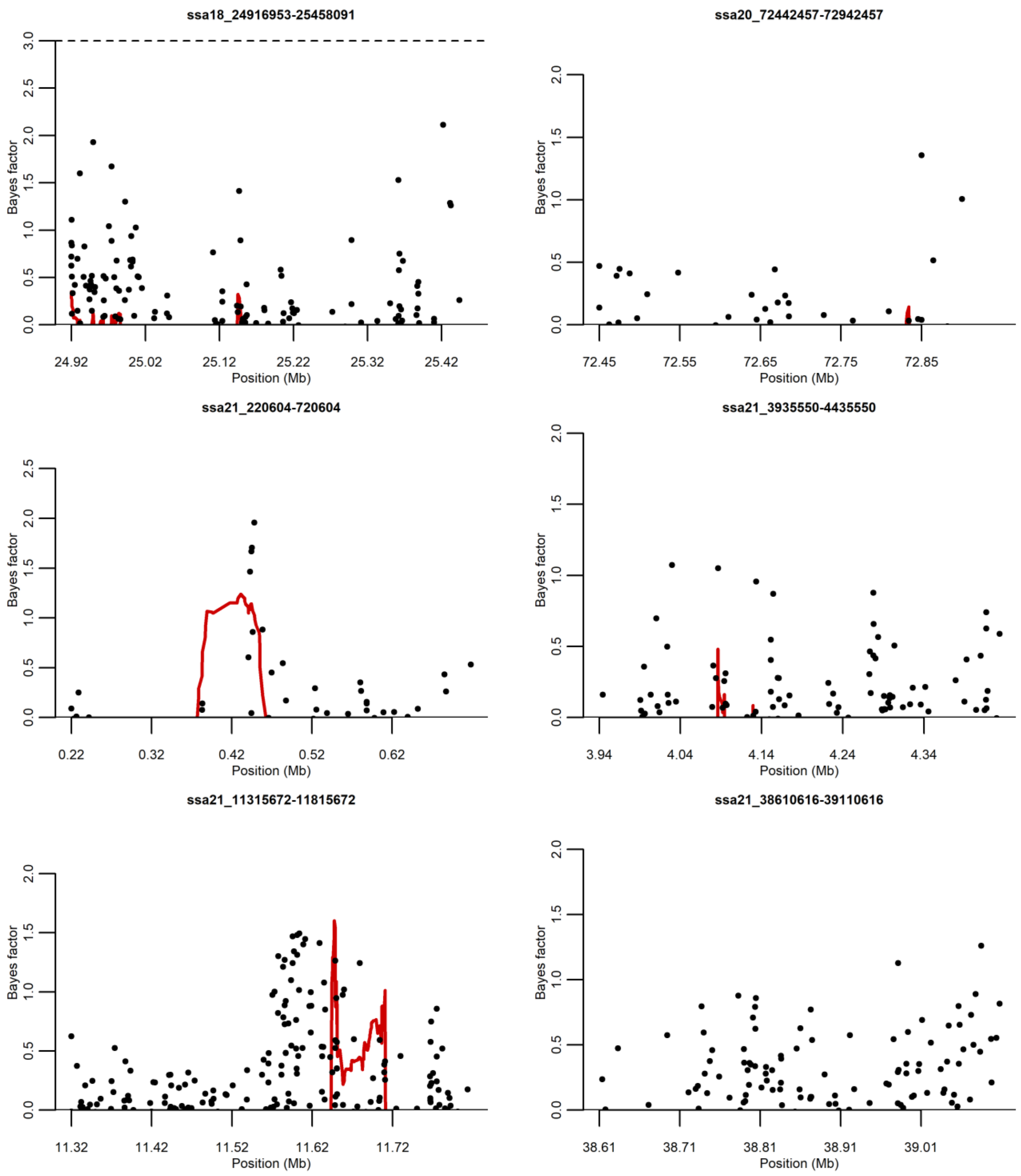
ssa18_24916953-25458091
Bayes factor
Position (Mb)
24.92
25.02
25.12
25.22
25.32
25.42
ssa20_72442457-72942457
Bayes factor
Position (Mb)
72.45
72.55
72.65
72.75
72.85
ssa21_220604-720604
Bayes factor
Position (Mb)
0.22
0.32
0.42
0.52
0.62
ssa21_3935550-4435550
Bayes factor
Position (Mb)
3.94
4.04
4.14
4.24
4.34
ssa21_11315672-11815672
Bayes factor
Position (Mb)
11.32
11.42
11.52
11.62
11.72
ssa21_38610616-39110616
Bayes factor
Position (Mb)
38.61
38.71
38.81
38.91
39.01

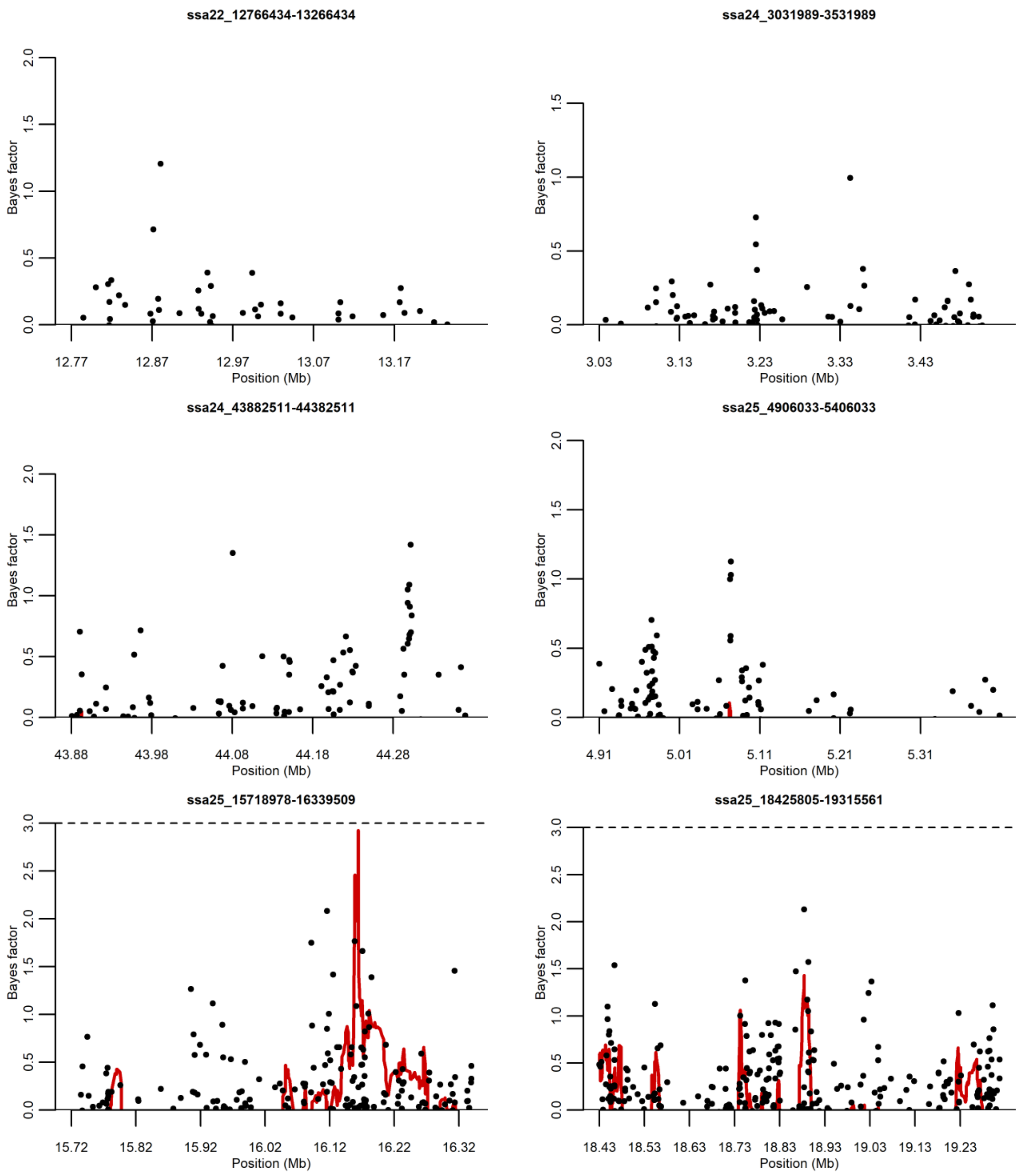
ssa22_12766434-13266434
Bayes factor
0.0
0.5
1.0
1.5
2.0
12.77
12.87
12.97
13.07
13.17
Position (Mb)
ssa24_3031989-3531989
Bayes factor
0.0
0.5
1.0
1.5
3.03
3.13
3.23
3.33
3.43
Position (Mb)
ssa24_43882511-44382511
Bayes factor
0.0
0.5
1.0
1.5
2.0
43.88
43.98
44.08
44.18
44.28
Position (Mb)
ssa25_4906033-5406033
Bayes factor
0.0
0.5
1.0
1.5
2.0
4.91
5.01
5.11
5.21
5.31
Position (Mb)
ssa25_15718978-16339509
Bayes factor
0.0
0.5
1.0
1.5
2.0
2.5
3.0
15.72
15.82
15.92
16.02
16.12
16.22
16.32
Position (Mb)
ssa25_18425805-19315561
Bayes factor
0.0
0.5
1.0
1.5
2.0
2.5
3.0
18.43
18.53
18.63
18.73
18.83
18.93
19.03
19.13
19.23
Position (Mb)

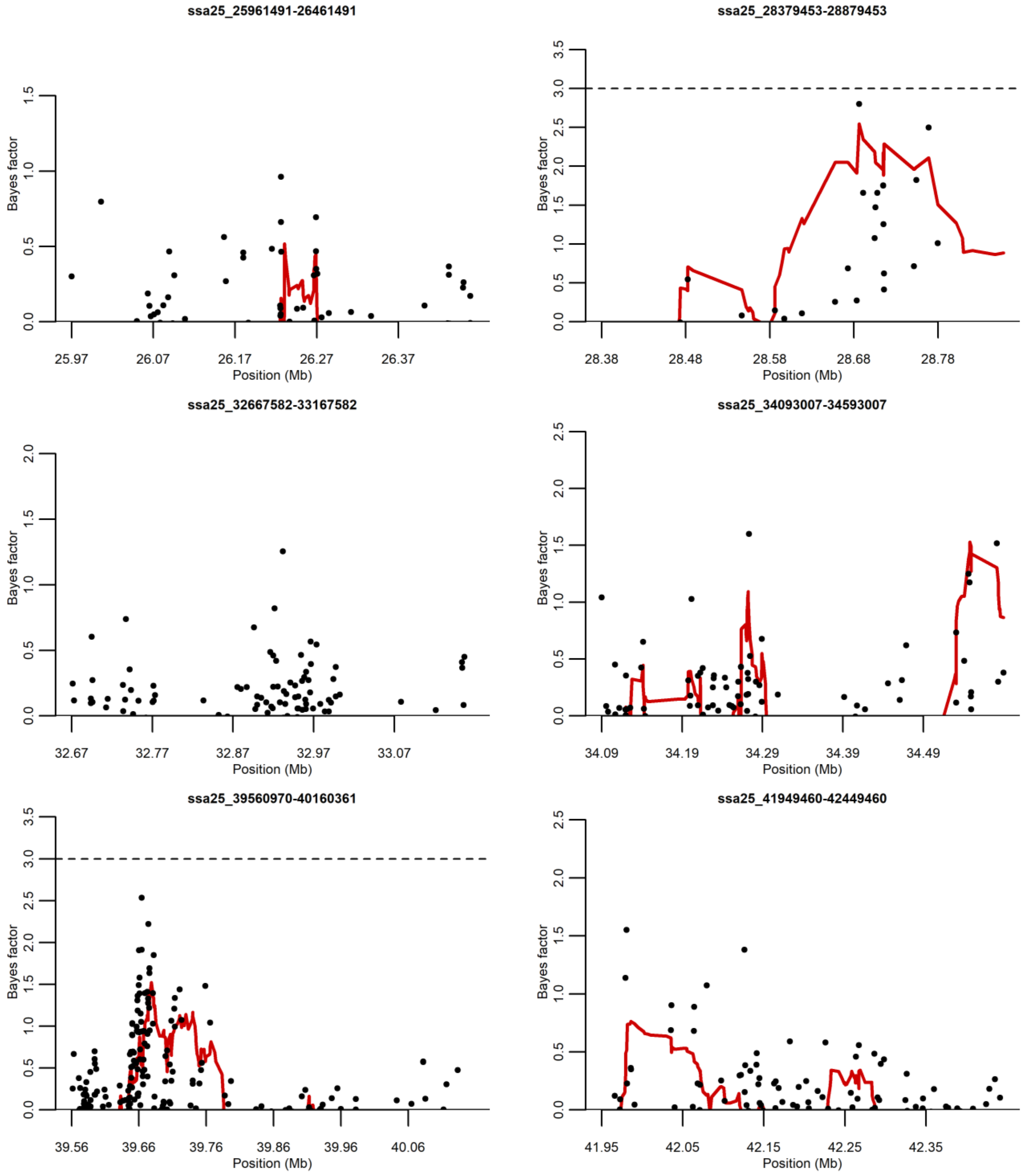
ssa25_25961491-26461491
ssa25_28379453-28879453
ssa25_32667582-33167582
ssa25_34093007-34593007
ssa25_39560970-40160361
ssa25_41949460-42449460
Bayes factor
Position (Mb)
25.97
26.07
26.17
26.27
26.37
28.38
28.48
28.58
28.68
28.78
32.67
32.77
32.87
32.97
33.07
34.09
34.19
34.29
34.39
34.49
39.56
39.66
39.76
39.86
39.96
40.06
41.95
42.05
42.15
42.25
42.35
0.0
0.5
1.0
1.5
2.0
2.5
3.0
3.5

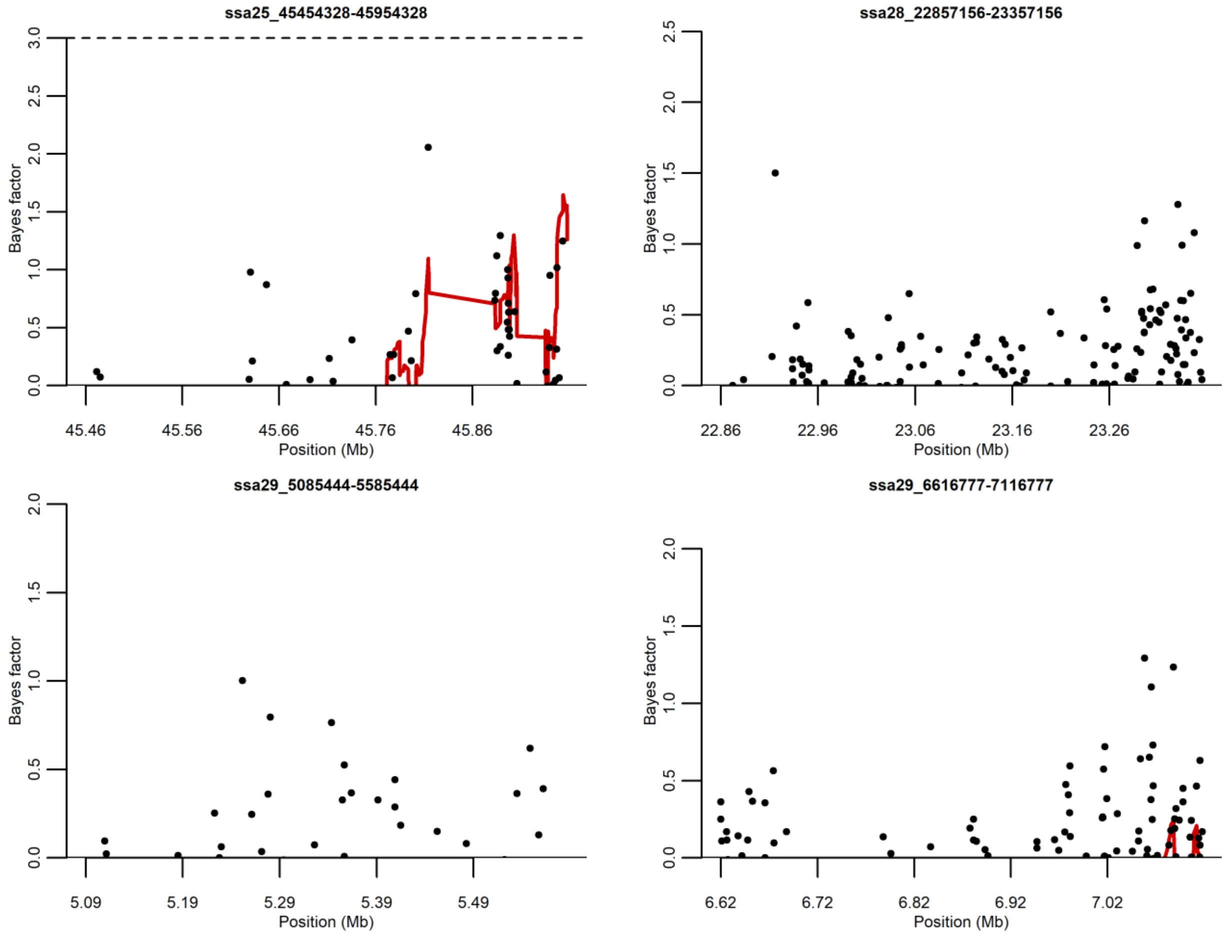


Supplementary Figure S2. Plots displaying single SNP associations (black points) and haplotype associations (red line) scores from *hapQTL* for each candidate region with Bayes factors less than 3. Y-axis shows the Bayes Factor indicating the degree of association strength. X-axis shows the position on the respective chromosomes.

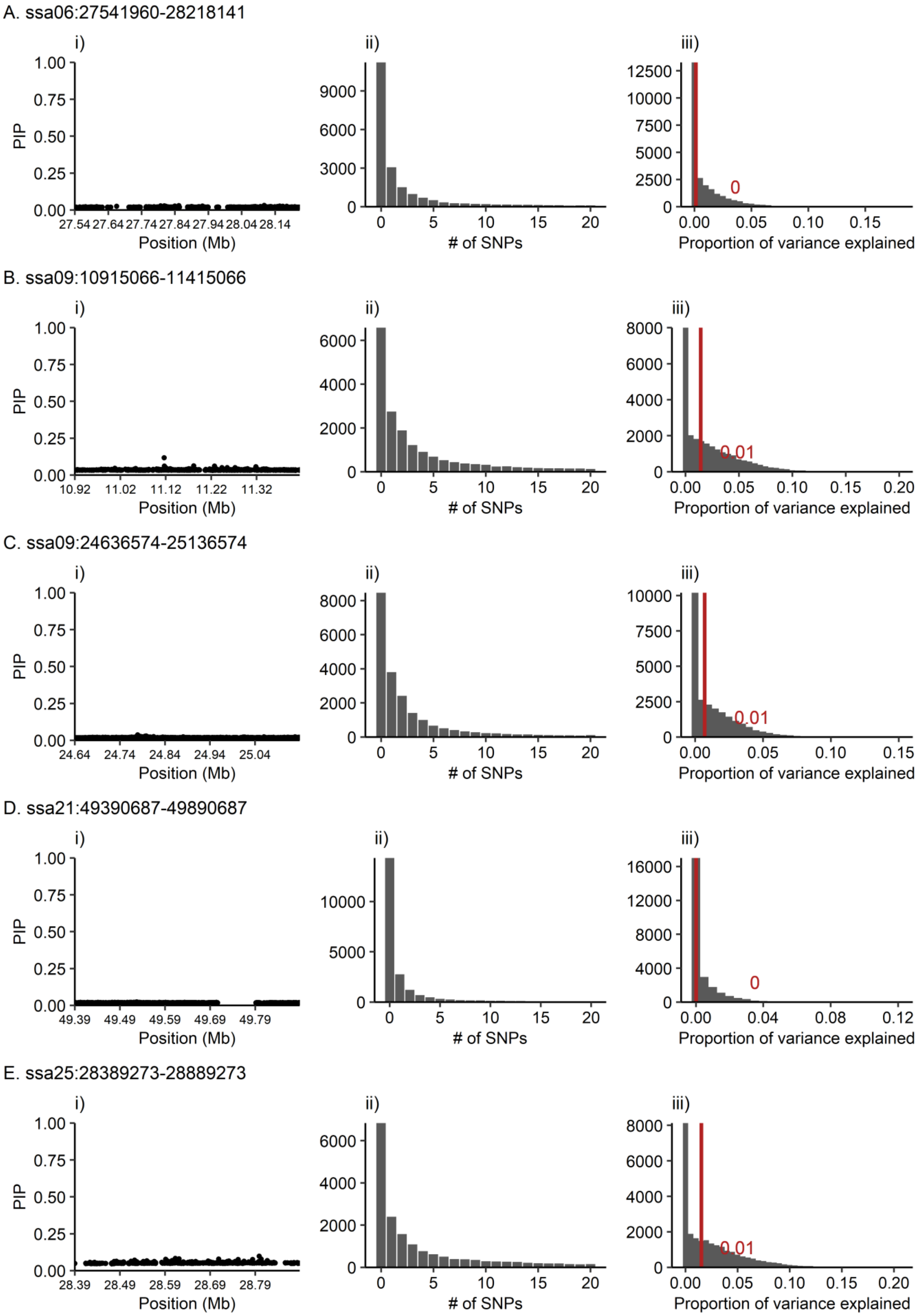
A. ssa06:27541960-28218141
i)
ii)
iii)
PIP
Position (Mb)
### of SNPs
Proportion of variance explained
0
B. ssa09:10915066-11415066
0.01
C. ssa09:24636574-25136574
0.01
D. ssa21:49390687-49890687
0
E. ssa25:28389273-28889273
0.01

Supplementary Figure S3. *PiMASS* results after regression of top-associated SNP identified in the initial *PiMASS* analysis. Each candidate region is shown as follows: A. ssa06:27541960-28218141, B. ssa09:10915066-11415066 C. ssa09:24636574-25136574, D. ssa21:49390687-49890687, and E. ssa25:28389273-28889273. Plots display the following results for each candidate region: i) posterior inclusion probability (PIP) indicating the probability of a SNP being included in a model explaining sea age at maturity variation, ii) truncated distribution of the number of SNPs included in a model explaining sea age at maturity variation per recorded iteration (2500) and iii) distribution of proportion of variance explained per recorded iteration (2500) where the red line indicates the median proportion of variance explained.

A. ssa06:27541960-28218141

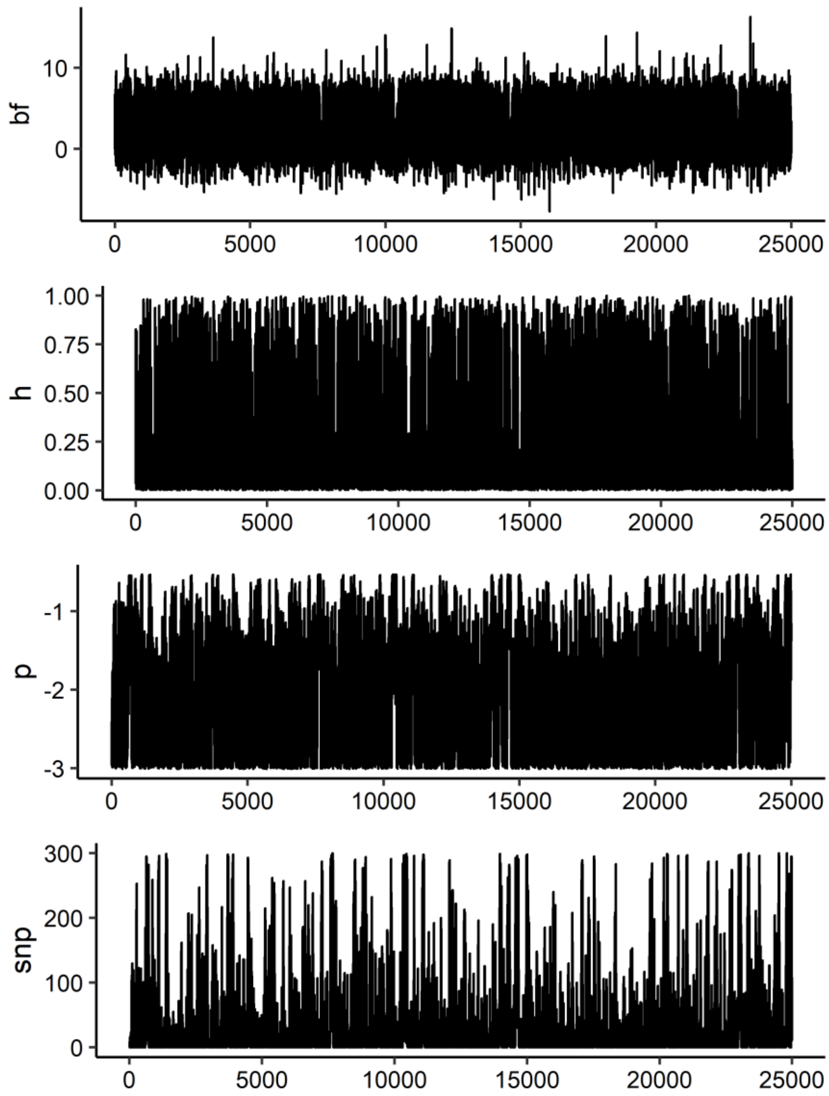

B. ssa09:10915066-11415066

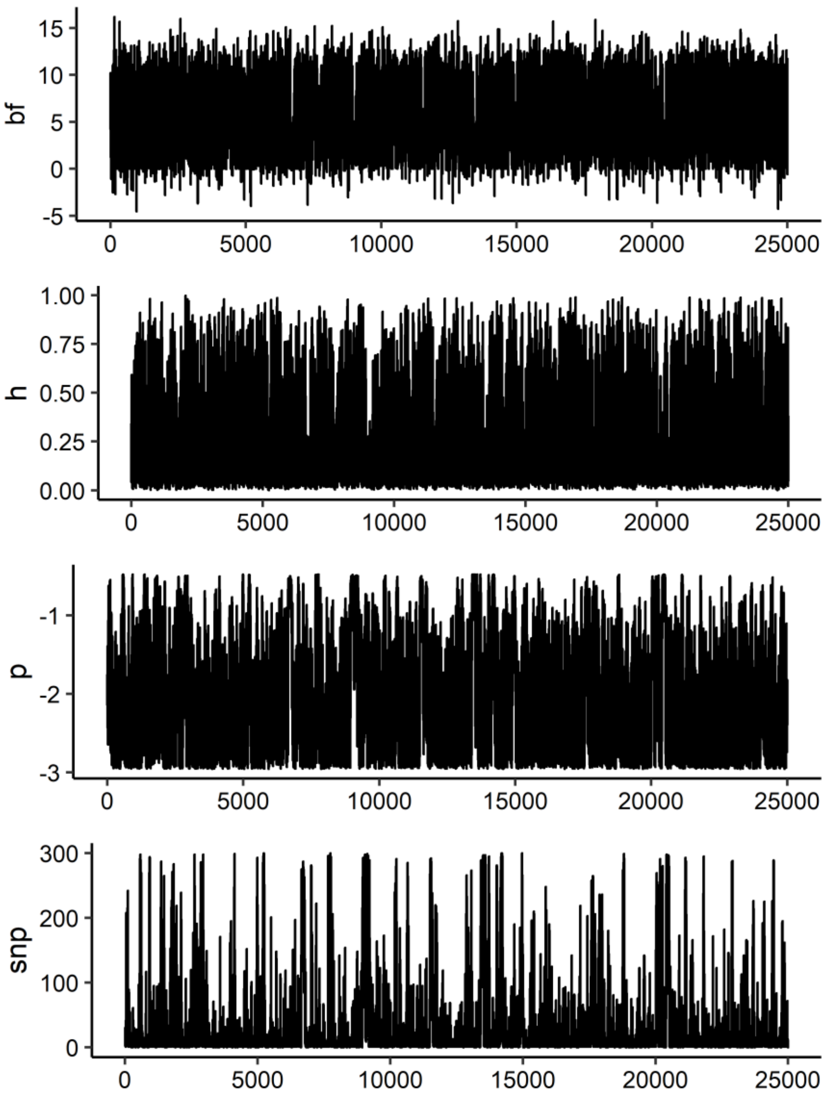

C. ssa09:24636574-25136574

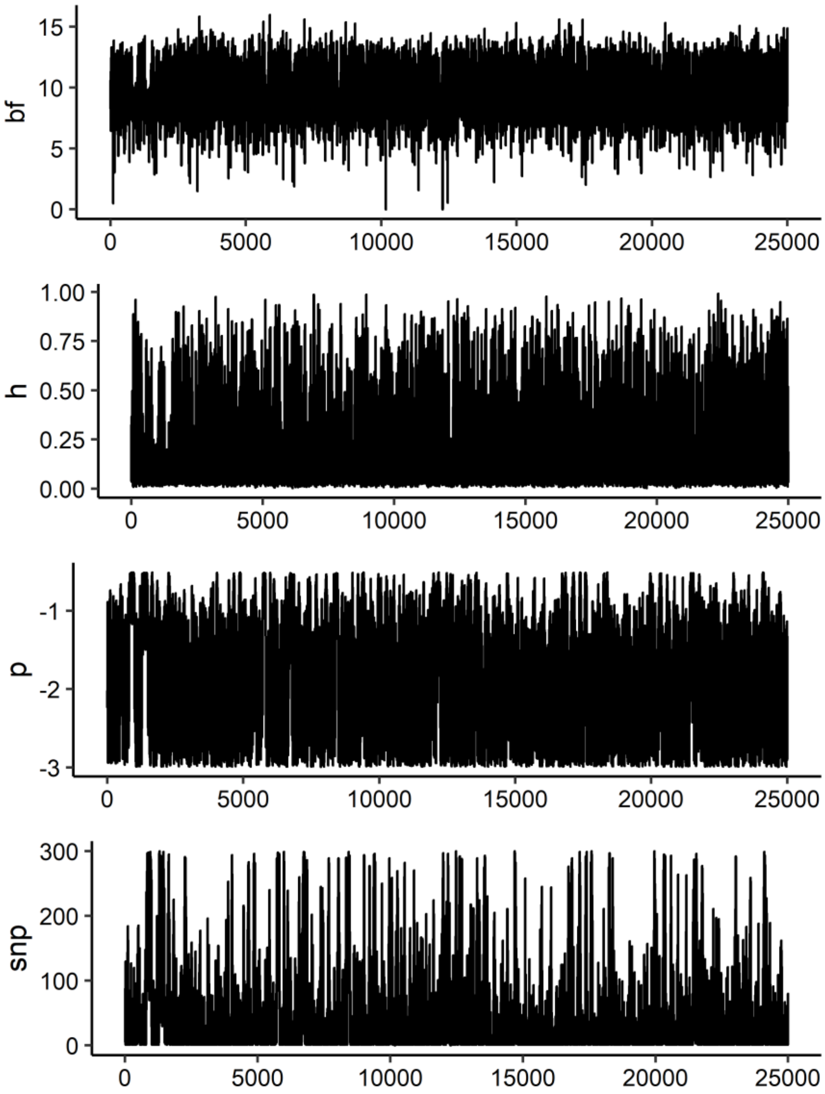

D. ssa21:49390687-49890687

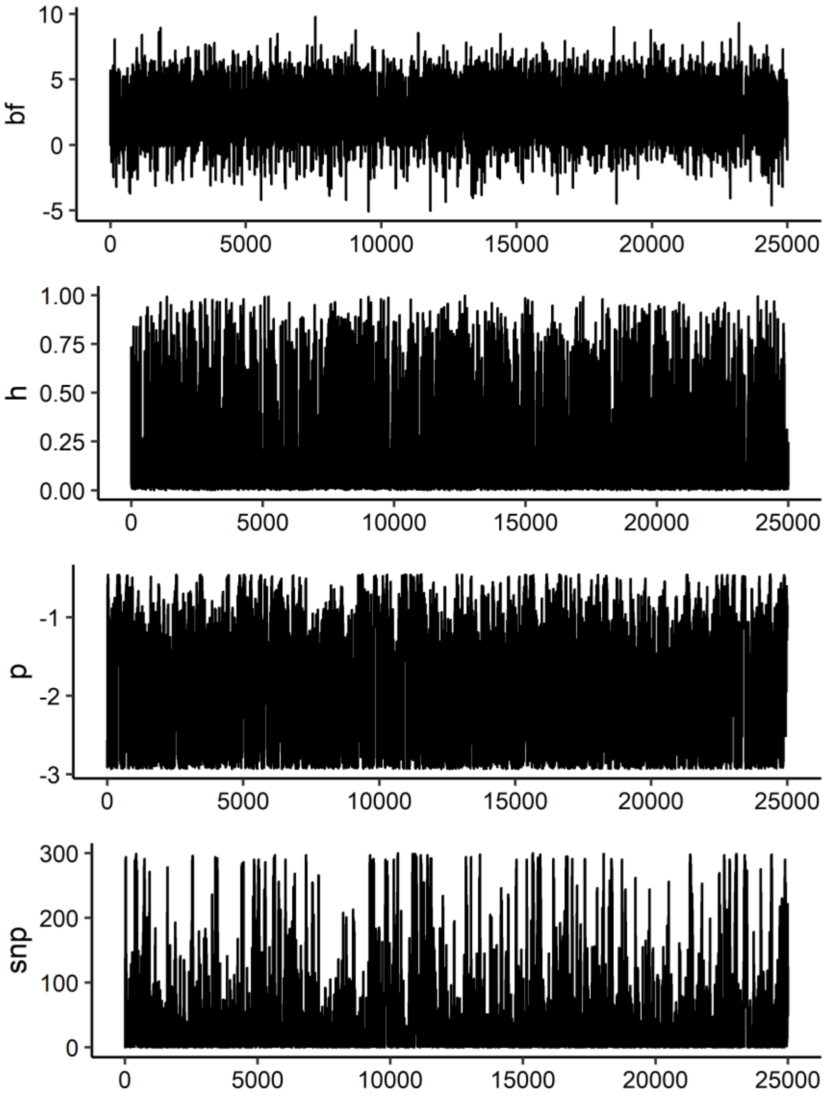

E. ssa25:28389273-28889273

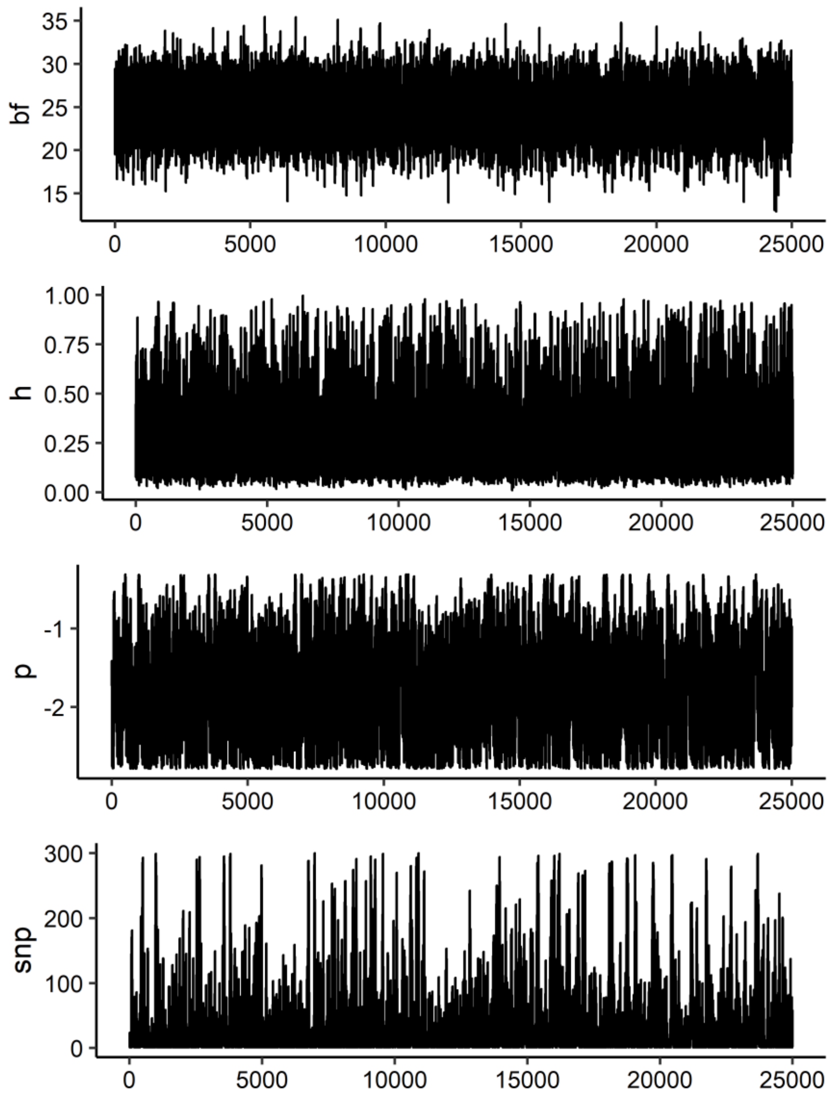


Supplementary Figure S4. Path of parameter values from *PiMASS* analyses for each candidate region: A. ssa06:27541960-28218141, B. ssa09:10915066-11415066 C. ssa09:24636574-25136574, D. ssa21:49390687-49890687, and E. ssa25_28389273-28889273. Values of log10 model probability (bf), heritability estimated with SNPs included in model (h), log10 prior probability of SNP being included in model (p) and number of SNPs included in model (snp), were recorded every 1000 steps.

A. ssa06:27541960-28218141

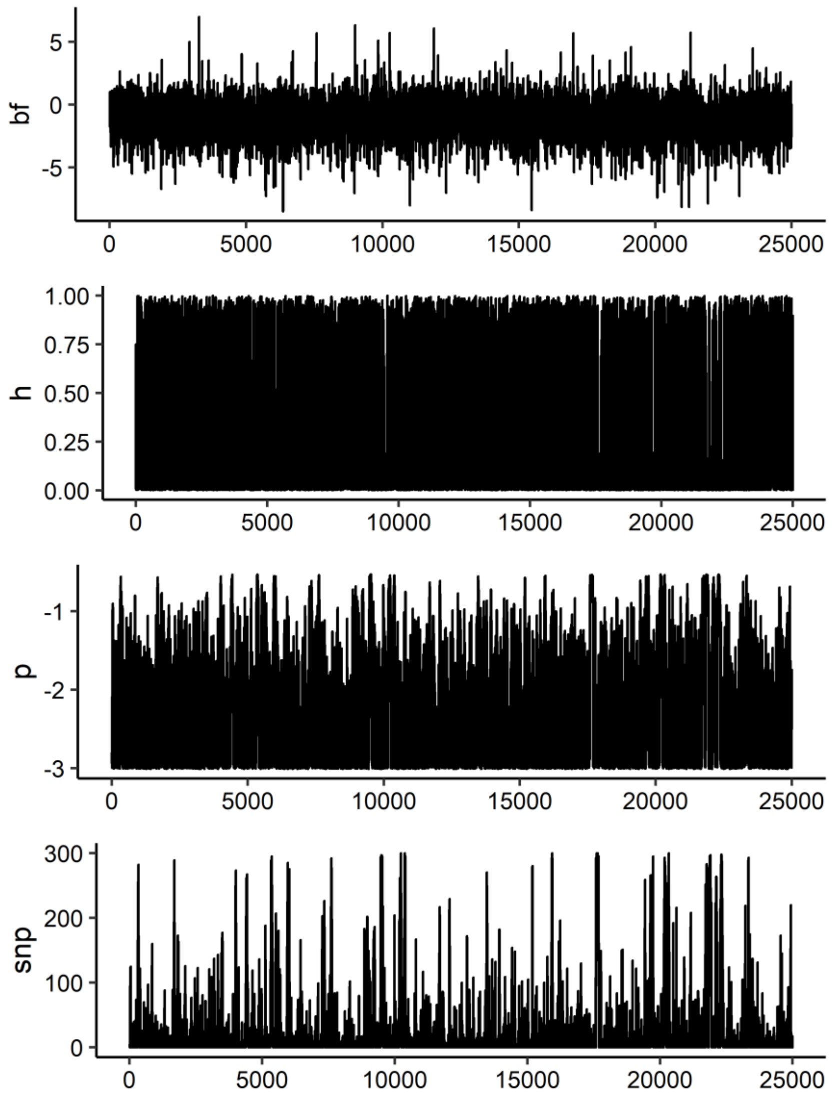

B. ssa09:10915066-11415066

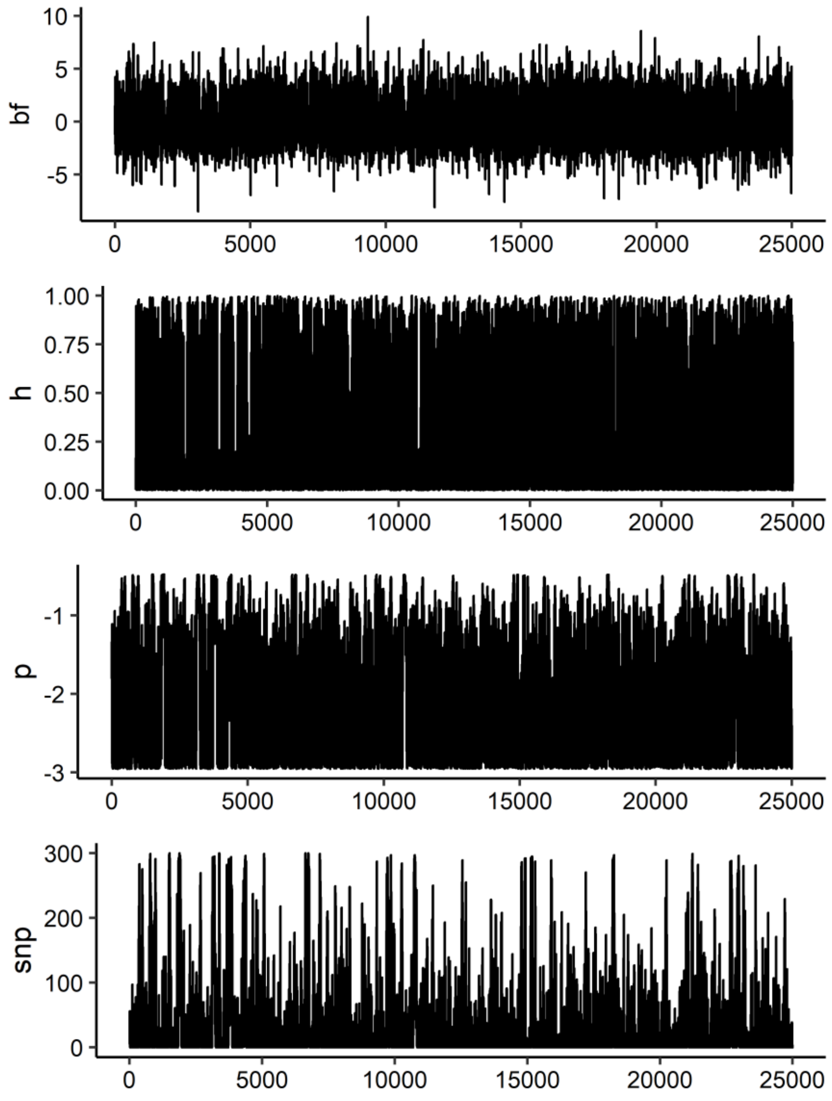

C. ssa09:24636574-25136574

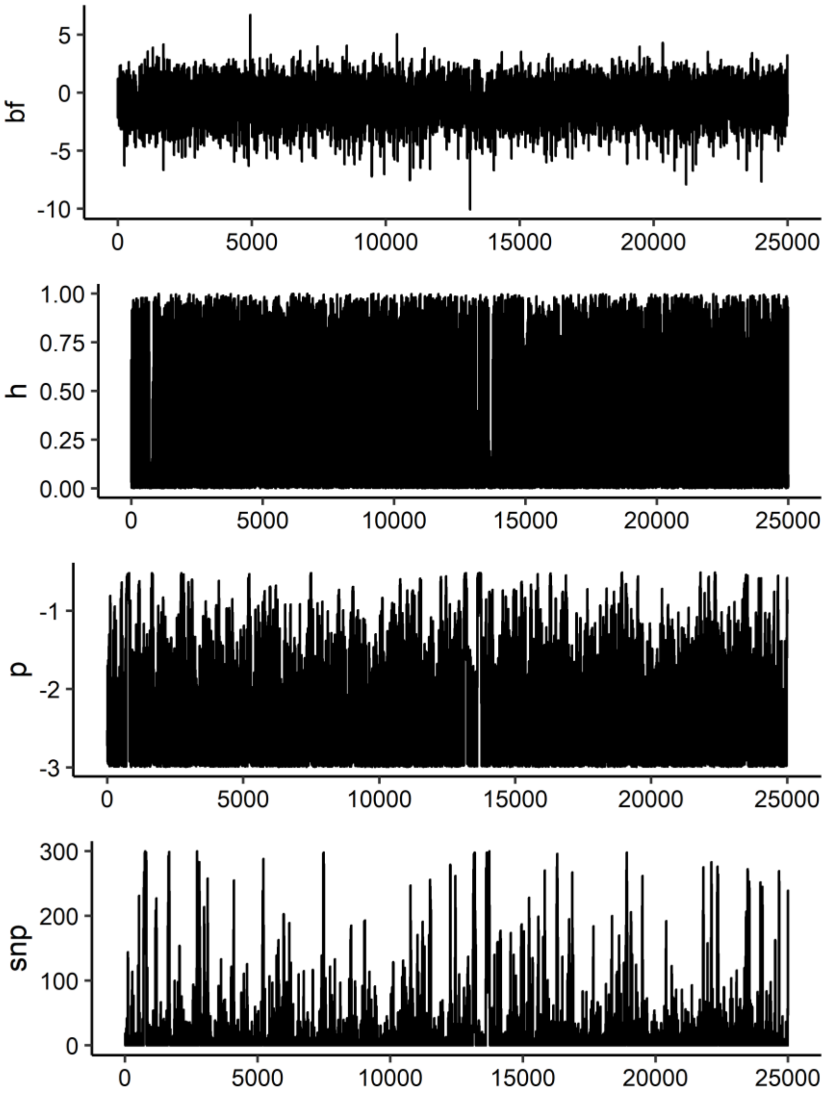

D. ssa21:49390687-49890687

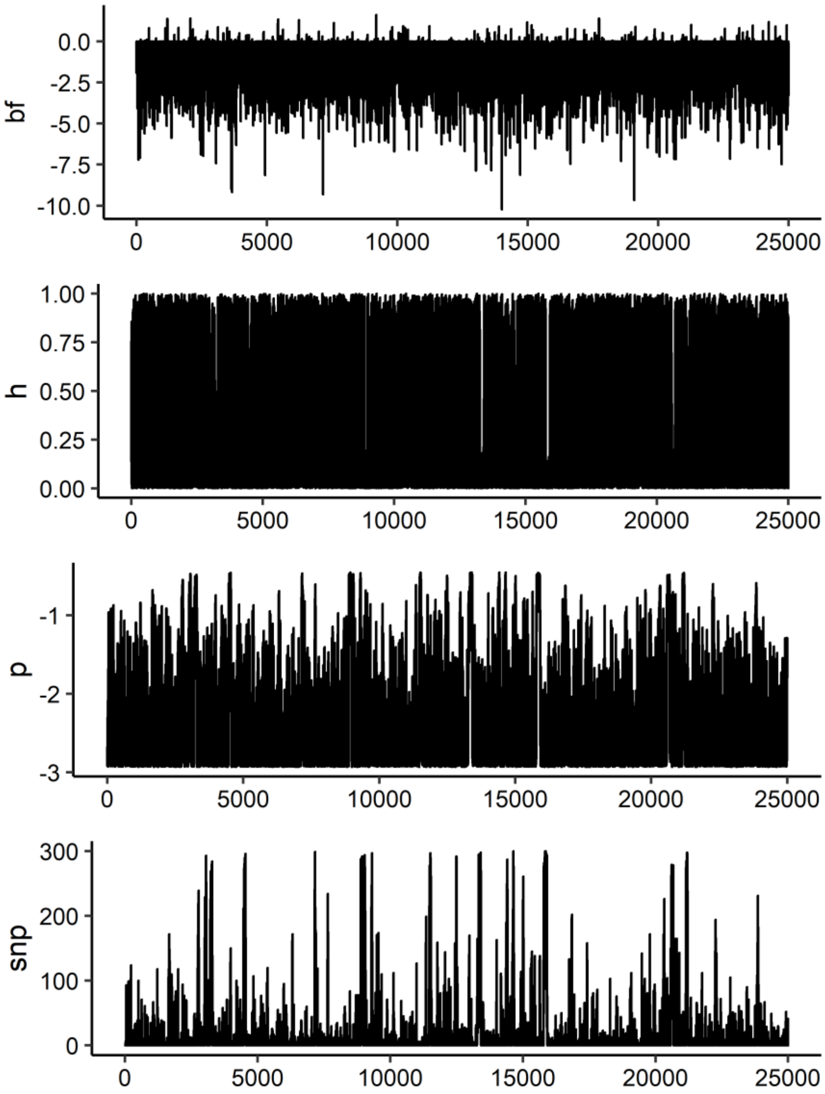

E. ssa25:28389273-28889273

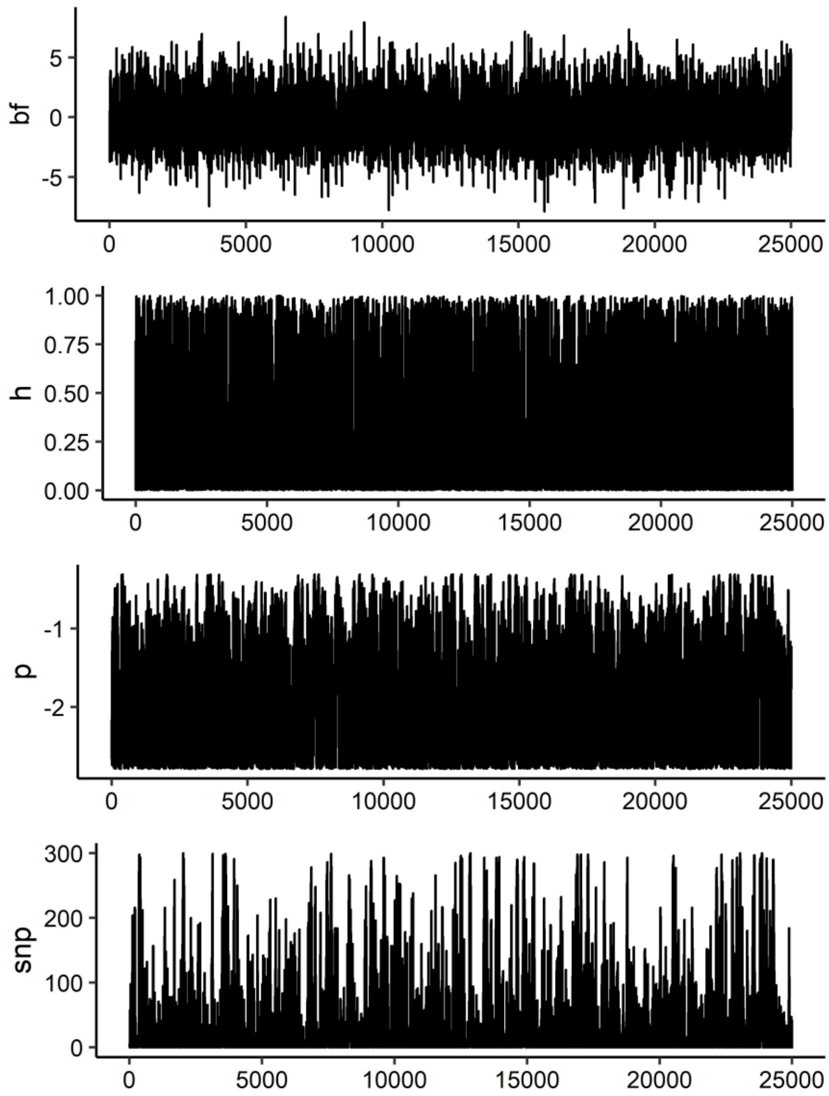


Supplementary Figure S5. Path of parameter values from *PiMASS* analyses for each candidate region following regression of top-associated SNP: A. ssa06:27541960-28218141, B. ssa09:10915066-11415066 C. ssa09:24636574-25136574, D. ssa21:49390687-49890687, and E. ssa25_28389273-28889273. Values of log10 model probability (bf), heritability estimated with SNPs included in model (h), log10 prior probability of SNP being included in model (p) and number of SNPs included in model (snp), were recorded every 1000 steps.
